# Stochastic choice drives variability in patch foraging decisions in humans and rats

**DOI:** 10.1101/2025.10.28.685063

**Authors:** Emma V. Scholey, Matthew A.J. Apps, Mark D. Humphries

## Abstract

Staying to exploit remaining resources or leaving to seek better options elsewhere is a fundamental decision across species. Optimal patch foraging theories propose deterministic rules for when to leave a depleting resource but real foragers show considerable variability in when they leave. Decisions between simultaneously-presented options are often assumed to follow a stochastic decision policy, adding randomness into the choice process to allow for exploration of potentially better alternatives. Whether a stochastic choice policy can account for variability in sequential foraging decisions, and what predictions such a policy makes for the mechanisms of foraging choice, are unknown. Here using patch foraging datasets in both humans (n = 39, n = 29) and rats (n = 8), we show that foragers making a stochastic choice of when to leave a patch is sufficient to explain their variability. We also show stochastic choice makes two unintuitive predictions, which we validate in our data. First, under a wide range of conditions, stochastic choice makes foragers’ leaving variability independent of the rewards available in the environment. Second, that foragers use a suboptimal internal function for setting their choice stochasticity from their environment’s average reward rate. Our findings suggest stochastic choice is an underappreciated but powerful contributor to foraging decisions, and highlight how behavioural variability, which is often overlooked, can reveal the algorithmic underpinning of decisions.

## Introduction

Neuroscientists have typically studied value-based decision-making as simultaneous choice between two or more rewarding options [1, 2, 3, 4, 5, 6, 7]. Yet, across species, real-world decisions often involve sequential encounters of a single option. Whether foraging for resources or deciding when to switch jobs, animals and humans must decide whether to stay and invest time on the current option, or forgo it in the hope that a better option is just around the corner [8, 9]. These kinds of ‘stay-or-leave’ decisions have a strong theoretical grounding in the field of behavioural ethology, where they are known as the ‘patch-foraging’ problem [10, 11, 12]. Neuroscience is increasingly turning to patch foraging to study the neural and cognitive processes underlying stay-or-leave decisions in health [13, 14, 15, 16, 17, 18] and psychopathology [19, 20].

In patch foraging, an animal harvests finite resources in a ‘patch’ and must decide whether to stay and spend time exploiting the remaining resources in the patch, or leave to explore the resources at other patches in the environment. Optimal behaviour in patch foraging is well defined by the marginal value theorem, [10, 12], which states an optimal forager should leave the current patch when its reward rate falls below the average reward rate in the environment. The marginal value theorem thus makes clear predictions about how patch and environment reward rates should influence leaving behaviour. Strikingly, many species follow these patterns of behaviour [11, 21, 22, 13, 16, 23]. However, since the marginal value theorem is a deterministic rule it predicts that a forager making a stay-or-leave decision in identical contexts should leave a patch at exactly the same time. Foragers do not: they display high intra-individual variability in their leaving choices [24, 25, 26], suggesting normative accounts of foraging omit key computational mechanisms essential to understand foraging decisions.

A simple explanation for this variability is that foragers use a stochastic decision rule rather than a deterministic one. Other areas of decision-making research assume this to be true of their subjects [27]. For example, reinforcement learning models of behaviour use a stochastic decision rule when selecting the next action from among multiple options, to allow exploration of potentially better alternatives than the action currently estimated to have the highest return [28, 3, 29, 30, 31]. Stochastic action selection can well capture human [3, 30, 32] and animal [33, 34, 35, 36, 32] behaviour on simultaneous choice tasks, where increased choice stochasticity leads to greater variability. Further, recent evidence shows that choice stochasticity adapts to environmental reward statistics in simultaneous choice tasks [37].

Whether these findings extend to sequential foraging decisions is unclear. Whilst foraging studies in psychology and neuroscience commonly model sequential foraging choices as stochastic [16, 38], it is unknown whether foragers’ choice variability is consistent with a stochastic decision rule. Confirming this hypothesis would thus provide support for widely-used assumptions in the literature; evidence against it would suggest a need to reevaluate the algorithmic basis for foraging decisions. If foraging choice is consistent with a stochastic decision rule, this would raise further important questions over the reasons for the stochasticity in sequential foraging decisions: whether it is just inherent, irreducible noise [39], deliberate noise enabling adaptation in volatile environments [3, 31], or an adaptive mechanism for producing unpredictability in behaviour, for example to evade predation [40]. Understanding the nature of a stochastic decision rule could also clarify if stochasticity constrains foraging behaviour, leading to compensatory and seemingly suboptimal foraging strategies such as overharvesting [24, 26].

Here, we test the hypothesis that a stochastic decision rule is sufficient to account for foragers’ intra-individual variability in patch leaving decisions. We consider a model with perfect reward information that includes reward-dependent and reward-independent contributors to the stochastic choice of when to leave a patch. As we show below, our model makes the unintuitive predictions that the variability of a forager’s leaving decisions does not depend on the reward available in their current environment or patch under a wide range of conditions, but does critically depend on how reward decays as a forager stays longer in a patch. Further, we show our model predicts that foragers have an internal function for weighting reward in the current patch by the average reward rate in the environment. We test these predictions by reanalysing data from three tasks in which foragers consistently make sub-optimal and variable patch-leaving decisions even in the same reward context, across different tasks and in both humans and rats.

## Methods

### Task descriptions

We analysed three existing patch foraging datasets. Two datasets used a human sample, and we refer to these as the ‘field-human’ and ‘berry-human’ tasks, representing the different games and resources collected. The third dataset used a rat sample, which we refer to throughout as the ‘rat’ dataset. Our study was not pre-registered. We conducted secondary data analysis in line with University of Nottingham policies that do not require ethical approval for such analysis.

#### Field-human

Data for the field-human task came from [18]. Forty healthy young adults (mean age = 24 yrs, range 20-30 yrs) were recruited, with one exclusion due to poor task engagement identified at debriefing (n = 39). Data on sex/gender, socioeconomic status, communities of descent, and race/ethnicity were not included. Participants completed a computer-based patch-leaving task, which was framed as a farm game. The goal was to collect as much milk as possible with a bonus financial remuneration based on the amount of milk collected. Participants encountered sequential patches with depleting milk and could decide at any point to leave the current patch and travel to the next one. The reward obtained so far in the patch was displayed as a continuously filling bucket, where the height of milk displayed was related to the decay rate of the patch, *g* (Table 1), and updated at a frequency of 20 Hz.

**Table 1:** Task parameter values.

|  |  | Field-human | Berry-human | Rat |
| --- | --- | --- | --- | --- |
| **Patch decay function** |  | Exponential | Exponential | Linear |
| **Patch decay rate, $g$, (ml)** |  | 0.075 | 0.11 | 0.008 |
| **Patch type** |  | low, mid, high | low, mid | low, mid, high |
| **Patch initial yields, $r_0$ (ml)** |  | 32.5, 45, 57.5 | 34.5, 57.5 | 0.06, 0.09, 0.12 |
| **Patch distribution** | Rich | 1/6, 1/3, 1/2 | 1/2, 1/2 | 1/3, 1/3, 1/3 |
|  | Poor | 1/2, 1/3, 1/6 | 1/2, 1/2 | 1/3, 1/3, 1/3 |
| **Travel times (s)** | Rich | 6 | 2.8 | 10 |
|  | Poor | 6 | 5 | 30 |

Patch reward decayed exponentially according to

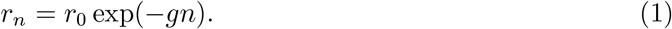

Participants were not explicitly told which patch type they were in (low, medium, high). Patch type was determined by the initial patch yield, with the same decay rate for all patches, meaning that a high yield patch would take longer to decay to the same reward rate as a low yield patch. When the participant decided to travel, they incurred a fixed travel time cost where no milk could be collected.

Participants completed two blocks of the task, in either a rich environment or poor environment, with a higher average reward rate in the rich environment. The average reward rate was manipulated by varying the patch distribution: participants were more likely to encounter a high yield patch in rich environments (Table 1). The type of patch encountered was pseudorandomised. Participants spent 10 minutes in each environment (block order was pseudorandomised), and the current environment was cued with a gold (rich) or green (poor) border around the screen. Participants were trained on the task with a practice session, where they encountered 10 patches in each environment.

#### Berry-human

Data for the berry-human task came from study two of [41]. Thirty-one healthy young adults were recruited, with two exclusions due to poor task engagement, identified as very short (*<*500ms) or very long average leaving times (*>*3 s.d. from sample average). This resulted in a final sample of 29 adults (mean age = 24.7 yrs, SD = 5 yrs, self-reported gender: 16 females, 13 males). Data on socioeconomic status, communities of descent, and race/ethnicity were not included. Participants completed a similar computer-based patch-leaving task to the farm game above, collecting berries instead of milk. Patch reward again decayed exponentially according to Eq 1. The main task differences are presented in Table 1 and summarised here.

Participants completed three blocks of the task, visiting four different farms in each block. The farms varied in whether they were rich or poor environments, but also whether the participant was collecting berries for themselves or for another person. We only include the ‘self’ blocks in our reanalysis of the data. Rich environments had a higher average reward rate than poor environments, determined by the travel time between patches, with a shorter travel time cost in rich environments (Table 1). There were two different patch types (low, medium), which were equally distributed in both environments. Participants spent 5 minutes in each environment (order was psedudorandomised), and the current environment was cued with a gold (rich) or green (poor) border when the participant first started the new block. Participants were trained on the task with a practice session for each environment.

#### Rat

Rat foraging data came from the travel time experiment of [23]. Eight adult Long-Evans rats were used. Details of the training and test schedule can be found in the original article.

For the travel time foraging task, rats decided whether to lever press to obtain 10% sucrose reward, or whether to nose poke to travel to a new, replenished patch. A cue light above the lever and inside the nose poke indicated the start of a trial when the rats could make a decision. A lever press (stay) decision resulted in reward in the reward magazine. To control for the reward rate within the patch, the next trial was initiated after an adjustable inter-trial interval (ITI), which depended on the decision time of the current trial, ensuring that all stay trials were equally long. On each successive stay decision, the rat received a smaller volume of reward, according to decay rate, *g* (Table 1) and the linear decay function

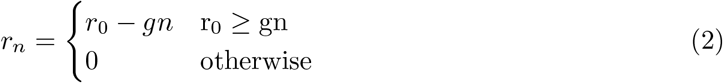

Here, *n* indexes each stay-or-leave choice. Patch type (low, medium, high) was determined by the initial patch yield, with the same decay rate for all patches. A nose poke (leave) decision resulted in the lever retracting for the length of the travel time. After this travel time, the other lever extended, and rats could again make a decision to lever press (stay) or nose poke (leave).

For different behavioural sessions, rats were tested in either rich or poor environments, determined by the travel time between patches. The three different patch types were equally distributed in both environments. For each environment, rats were trained for 5 sessions across consecutive days followed by 5 testing days. The order of environments was counterbalanced. Rats visited a range of 87-236 patches per environment.

### Behavioural analysis

Analysis of behavioural data was run in MATLAB 2023b, R v4.4.1 with RStudio, and JASP 0.19.3. All significance tests were two-tailed with a threshold of 0.05. Within each dataset, repeated measures were taken from the same sample. Correlations of model parameters with subject patch leaving times were calculated with Spearman’s rank correlation coefficient in MATLAB. For comparisons of variability (*SD*_*leave*_) in early versus late phases, and for comparisons of *B* in rich versus poor environments, normally distributed data were analysed using Student’s t-test (*ttest* in MATLAB), and a Wilcoxon signed rank test was used as a nonparametric alternative (*signrank* in MATLAB). For the rat task, we used Wilcoxon signed rank test by default. Confidence intervals for the signed rank test were calculated using bootstrapped 95% CIs on the median paired difference with 5000 samples. Effect sizes are reported using Cohen’s *d* for parametric tests, and robust Cohen’s *d* for non-parametric equivalents, using the *meanEffectSize* function in MATLAB. We computed Bayes factors for the null hypothesis (*BF*_01_) using either a Bayesian t-test, or a Bayesian Wilcoxon signed-rank test in JASP 0.19.3 with 1000 samples for Markov Chain Monte Carlo (MCMC) simulations (requiring a Gelman-Rubin statistic less than 1.05 to indicate convergence). We used the default Cauchy prior with a scale of 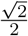. The Bayes factor *BF*_01_ represents how many times more likely the data are under a null hypothesis of no effect compared to the alternative hypothesis that there is an effect. Traditionally, a *BF*_01_ larger than 3 (equivalent to *BF*_10_ smaller than 1/3) is considered moderate evidence in favour of the null hypothesis [42]. For all statistical analyses, we confirmed that the data met the assumptions of the tests used, using a combination of visual inspection and testing for a normal distribution using the Shapiro-Wilk test [43].

### Optimal predictions for leaving time

The marginal value theorem predicts that an optimal forager should leave a patch when the patch reward rate *r* at continuous time *t* in the patch falls below the average reward rate *ρ* in the environment [10]. Thus the leaving rule is formalised as:

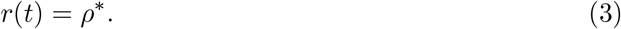

The optimal leaving times *t*^∗^ predicted by the marginal value theorem were calculated by substituting the equation for patch reward decay on the left side of the leaving rule, and the optimum average reward rate on the right side, and solving for *t*.

For the tasks with exponential decay of patches (given by Eq 1), optimal leaving times were thus defined as:

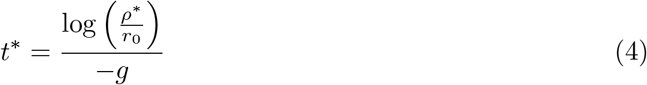

For the tasks with linear decay of patches (given by Eq 2), optimal leaving times were defined as:

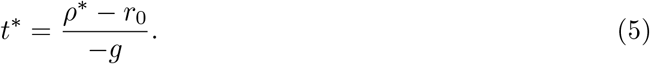

The initial reward rate *r*_0_ and decay rate *g* were already defined. We calculated the optimum average reward rate *ρ*^∗^ by simulating a foraging agent visiting sequential patches with task parameters defined in Table 1. We set *ρ*^∗^ at a fixed value and computed the optimal leaving time *t*^∗^ for each patch using Eq 4 (exponential) or Eq 5 (linear). Based on these patch encounters and leaving times, we calculated the accumulated total reward and total time (accounting for travel times) until the total block time passed. The observed average reward rate was computed as 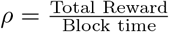 and compared to the simulated optimal average reward rate *ρ*^∗^. This process was repeated iteratively until the observed *ρ* converged on the simulated *ρ* (with a tolerance of 0.0001). The initial value of *ρ*^∗^ was set to 30 (units/s) and each new iteration used the observed *ρ* estimate from the previous iteration. The maximum iterations were set to 1000.

### Model analysis

Our stochastic stay-or-leave model was of the general form

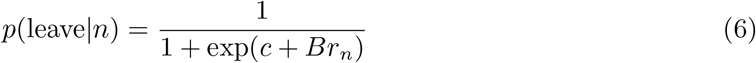

which calculates the forager’s probability of leaving a patch in time step *n* given the current reward obtained in the patch, *r*_*n*_. This model includes both reward-independent (*c*) and reward-dependent (*B*) contributions to stochastic choice.

To generate *a priori* predictions of our model in Eq 6 for Figure 4 and Supplementary Figures 2 and 5, we calculated the expectation and variance for its leaving times. We used the standard equations for expectation, *E*(*x*) = ∑*xp*(*x*), and variance, *V ar*(*x*) = ∑ (*x* − *E*(*x*))^2^*p*(*x*), of a random variable *x*. In the patch foraging tasks, the random variable is *n*, the number of elapsed timesteps.

The expected leaving time is therefore

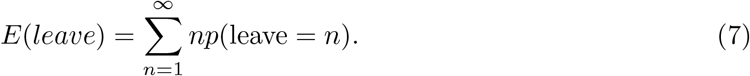

Similarly, the variance in leaving time is therefore

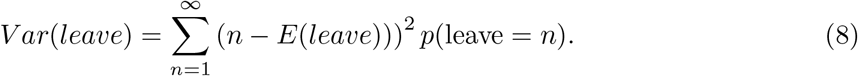

The probability *p*(leave = *n*) is the probability that timestep *n* is the first timestep at which leaving occurs. Each timestep *n* has only two possible outcomes (stay or leave); the sequence of timesteps *n* is a Bernoulli process. The probability is thus

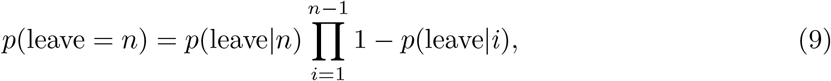

where the first term is the probability of leaving on this timestep, *n*, and the second term is the cumulative probability of not having left on all the previous timesteps.

To generate *E*(*leave*) and *V ar*(*leave*) for each task, we calculated *p*(leave = *n*) using Eq 9 and our initial model in Eq 6. The reward *r*_*n*_ at each timestep was determined by the patch reward parameters in Table 1 and the reward decay given by Eq 1 (exponential decay for human tasks) or Eq 2 (linear decay for the rat task). We could then calculate *E*(*leave*) and *V ar*(*leave*) for each task, over a range of *B* or *c* values. To compare variability predictions to data we plotted 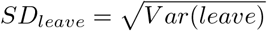.

To generate specific predictions for *E*(*leave*) for each task in Supplementary Figure 2, we assumed each timestep *n* to be 1 second (human tasks) or one harvest (rat task), and found the values of *B* or *c* that approximately reproduced the mean leaving times of subjects. We did this by finding the value of *B* or *c* that reproduces the average group leaving time in each environment for the mid-yield patch only, since, for all tasks, the likelihood of encountering this patch was matched between environments. We then used this same value of *B* or *c* to calculate the expected leaving time for low and high-yield patches in each environment. We took the same approach to generate the model predictions for optimal leaving times, as defined by the marginal value theorem, instead matching *B* or *c* to the optimal leaving times rather than subject leaving times.

### Model fitting

We conducted a two-step model fitting pipeline. In a first step, we compared variants of our stochastic models in which either the reward-independent bias, *c*, the reward-weighting parameter, *B*, or both varied between environments, in order to identify which mechanism best accounted for environment and patch effects on mean leaving times and *SD*_*leave*_ while keeping the model space tractable. We also tested two control models, a ‘vary *B*’ model that varied *B* between environments with no reward-independent bias (*c* = 0); and a ‘fix *B*, fix *c*’ model that fixed both parameters between environments. We did not test a ‘vary *c*’ variant with no reward-dependent weighting, as our analysis of Eq 6 showed this produces pathological leaving behaviour with the same expected leaving time for all patch types.

For all models, we assumed that *r*_*n*_ was known to the subject. A model where *r*_*n*_ was also estimated via a delta rule at each timestep did not change model fits; these results are not shown. For model fitting, *B* was constrained between 0 and 50, and *c* between -60 and 10.

We fit our models to subject data for each task using maximum likelihood estimation (MLE) i.e., minimising the negative log likelihood of the data given the model parameters. For each subject, gradient descent was performed with *fmincon* in MATLAB 2023b. The starting parameters were randomised to avoid local minima, with 50 random parameter seeds constrained within parameter bounds. To obtain the best fit parameters, we selected the fitting iteration with the lowest negative log likelihood. For model comparison at the group level, we calculated the summed Bayes Information Criterion (BIC) [44] across subjects.

For fitting the model to the human tasks, since the patch leaving decisions were continuous, we converted them to *N* timesteps by rounding the leaving time for each patch to the nearest second, and set the *N* of each patch equivalent to the number of seconds, as we did for the model analysis above. For the rat task, the stay-or-leave decision was already discretised.

#### Model and parameter recovery

We performed model and parameter recovery simulations to check the validity of the model comparison and parameter estimation [45]. For model recovery, we simulated 100 agents for each model displayed in Figure 5, drawing random parameters for each simulation from a uniform distribution based on subjects’ fitted parameter estimates for that model. For *B*, we bounded the distribution between 0 and the mean plus 3*×* standard deviation of *B*. For *c*, we bounded the distribution between the mean ± 3*×* standard deviation of *B*. All models were then fit separately to the simulated data from each model, using the MLE with 5 random initial parameter seeds. We took the iteration that produced the lowest negative log likelihood, calculated the BIC for each model, and approximated posterior probabilities [46]. We converted these into exceedance probabilities using the ‘spm BMS’ function from SPM25 to create a confusion matrix of model exceedance probabilities. This matrix indicates the confidence that simulated data from each model is best fit by the same model, where an identity matrix indicates good model recovery.

For parameter recovery, we simulated a further 100 agents for the winning model, drawing parameters as for model recovery. We fit the model to the simulated data using MLE, with 5 random initial parameter seeds to avoid local minima, taking the parameters that produced the lowest negative log likelihood. We report the Spearman’s correlations between simulated and recovered parameters, with all parameters showing excellent recovery (*r*_*s*_ ≥ 0.87 for all tasks).

#### Models for *B*

In our second model fitting step, we carried forward the winning model to compare internal functions that could map average reward rate onto the reward weighting parameter, *B*. Since our model fitting identified that ‘*vary c*’ models were a poor fit to the data, and thus *c* was approximately constant regardless of the average reward rate, we did not include functions linking *c* to average reward.

To determine whether we could derive *B* as a function of each subject’s experienced average reward rate, we fit four different models

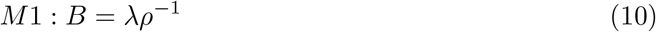

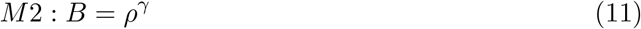

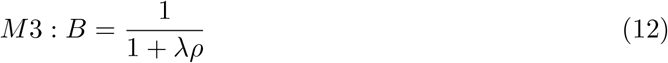

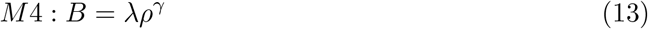

Each model is formulated so that lower average reward rates are translated into higher *B*, and thus increased reward weighting of stochastic choice. These models represent different simple functional forms of how *ρ* is transformed to *B*; they do not make strong theoretical or biological claims about this transformation.

We computed each subject’s experienced average reward rate, *ρ*, separately in rich and poor environments, by taking the mean of the average reward rate across blocks of the same environment. This is an omniscient model, where the agent knows a fixed estimate of *ρ*. For model fitting, *λ* was constrained between 0 and ∞, and *γ* between -5 and 5.

#### Learning models

We considered that whilst the subjects in our tasks were well-practised and thus we assumed a stable estimate of the average reward rate, typically the environment’s statistics are not initially known to the forager and must be learned through experience.

We extended the model defined by Eq 6 to include a standard delta rule, inspired by previous models [16]. This model estimates the average reward rate, *ρ*_*n*_, given the reward *r*_*n*_ at each timestep *n*, with learning rate, *α*:

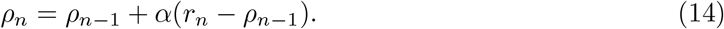

Then, the foraging agent updates *B*_*n*_ according to

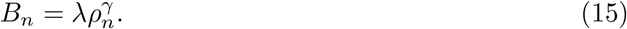

The agent then decides whether to stay or leave after timestep *n* according to Eq 6.

This model accounts for the passage of time both within the patch and spent travelling between patches. For each timestep during travel the agent receives no reward (*r*_*n*_ = 0) and thus the estimated average reward rate, *ρ*, was lower in environments with longer travel times (such as in the berry-human and rat tasks), and in environments with poorer patch yields (field-human task), in turn leading to higher reward-weighting (*B*) of stochastic choice in poor environments. For the berry-human task, the travel time in rich environments was 2.8s, which we rounded to 3s so that each timestep was equivalent to 1 second.

Taking a similar approach to previous studies [16], we initialised *ρ* to the average of the optimal average reward rates across both environments. In each subsequent environment, *ρ* was carried over from the previous environment. Learning rate, *α*, was constrained between 0 and 1 for model fitting.

We also fitted and compared learning versions of all potential models translating *ρ* into *B* (M1-4). We found that learning models of the general form of M4 were still the best fitting models overall and were the best model for the majority of subjects across all tasks, either when *γ* was fixed (*γ* = −1) or variable across subjects.

### Model simulations

To check the qualitative fit of our models against the real leaving choices of subjects, we ran simulations for each of the winning models. In each iteration (n = 50), we simulated each subject with their best-fit parameters from the winning model. We simulated the sequential encounters of patches and their stochastic stay-leave decision at each timestep, based on these parameters. For each virtual subject, we set their block order and experienced average reward rate as equivalent to the real data. We then calculated the average leaving time and standard deviation of the virtual subject for each patch type and environment. We averaged leaving times and standard deviation in leaving times over iterations.

## Results

### Humans and rats make stochastic leaving choices

We examined the variability of patch leaving across three published patch foraging tasks involving two species (Figure 1a; Table 1 in Methods). In the field-human (N=39) [18] and berry-human tasks (N=29) [41], healthy human subjects completed a computerised task to maximise the amount of milk or berries collected within a time limit. Subjects could leave their current patch at any time and travel, emulated by a time-out screen, before arriving at a new, replenished patch. In the rat task (N=8) [23], rats harvested depleting sucrose reward by lever pressing, and made decisions to harvest or leave at discrete decision points; leaving was signalled by a nose-poke, and travel between patches was emulated by a time-out before the new patch was available for harvesting using the lever. Thus leaving behaviour in the human tasks was measured as the patch leaving time in seconds, whereas in the rat task it was the number of harvest decisions per patch.

**Figure 1:**
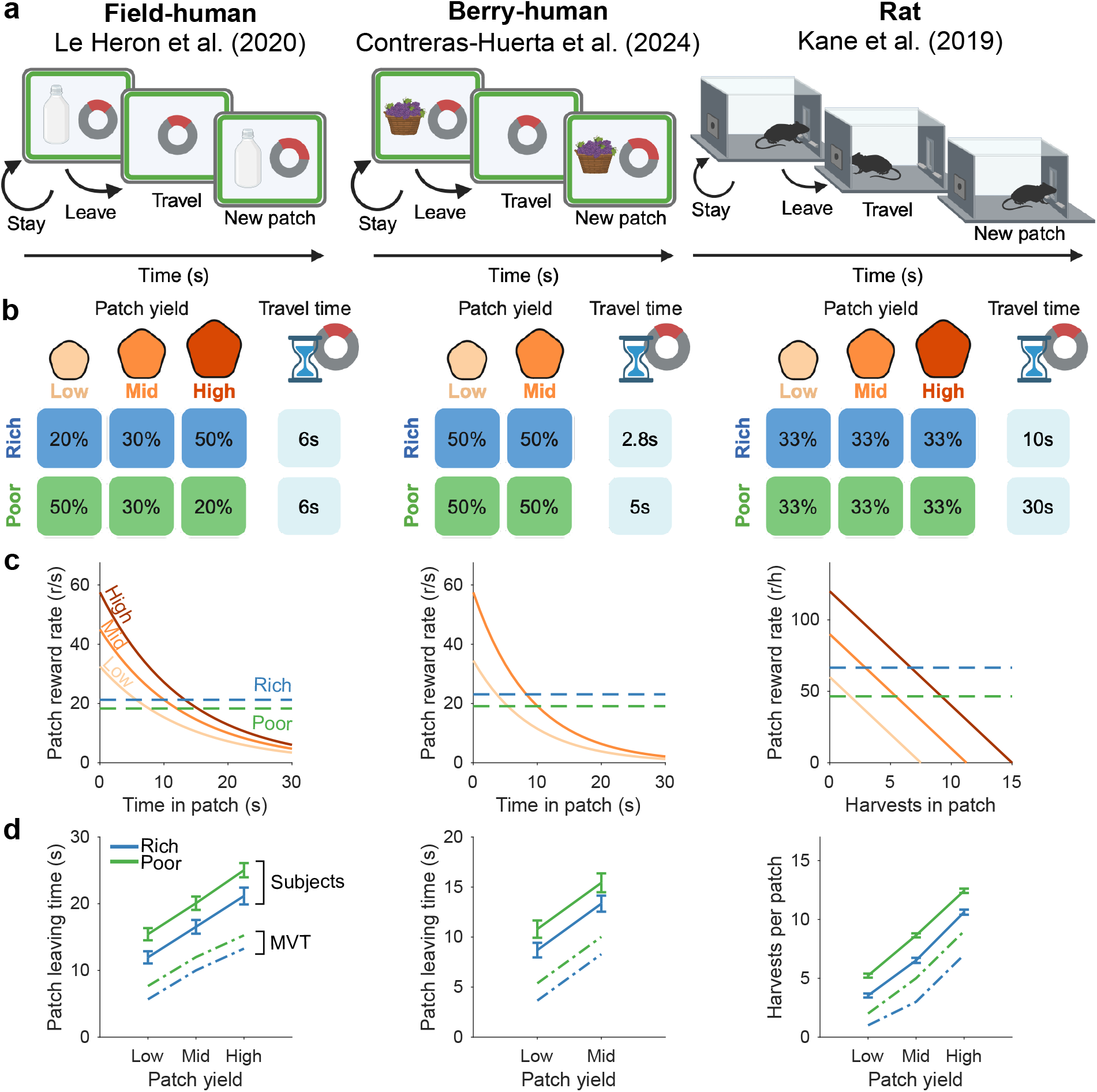
Consistent foraging behaviour across tasks and species. **a**: Foraging tasks. In the field-human and berry-human tasks, humans performed a computerised game collecting resources continuously delivered in patches, and decided when to leave to travel to a new patch. In the third task, rats could harvest sucrose water from the patch by lever-pressing. Each subsequent lever press resulted in depleting sucrose reward. Instead of lever-pressing, rats could instead nose-poke in the rear port, to replenish reward at a new patch after a travel time delay. **b**: Task parameters. All tasks have different patch types distinguished by their initial yield. All tasks presented environments in separate blocks of patches. The proportions of each patch type encountered in each environment are shown in the coloured boxes. **c**: Task reward rates. The patch reward rate decayed exponentially for all patches (solid lines) in the human tasks, and linearly in the rat task. Human tasks shows reward per second (s), whereas rat task shows reward per harvest (h). Average reward rates (dashed line) were higher in rich than poor environments. **d**: Group mean patch leaving decisions (solid line) compared to optimal behaviour (dash-dot line), for each patch and environment. Error bars show one SEM. Field-human: n = 39, berry-human: n = 29, rat: n = 8. Panels a and b created in BioRender. Scholey, E. (2026): https://BioRender.com/rjtixnl

Manipulations of patch and average reward rates in these tasks were inspired by real-world foraging environments. Subjects encountered different patch types distinguished by their initial yield (Figure 1b), which then decayed exponentially in the human tasks or linearly in the rat task (Figure 1c). Patches were encountered in either rich or poor environments (Figure 1b). Rich environments had a higher average reward rate than poor environments, but the average reward rate was manipulated in different ways in each task: by the likelihood of encountering high versus low yield patches (field-human), or the duration of travel times between patches (berry-human, rat). The rat dataset had a larger difference in travel times between environments, and thus a larger difference in average reward rates (Figure 1c).

We first reanalysed these data to confirm that, as reported in the original studies, both humans and rats showed patterns of patch-leaving behaviour predicted by the marginal value theorem (Figure 1d, Supplementary Note 1, Supplementary Table 1). Subjects in all tasks showed a patch effect, harvesting high yielding patches for longer than low yielding patches. Subjects also showed an environment effect, where they harvested patches of the same yield for longer in a poor environment than in a rich environment. In all tasks subjects also showed the often-observed overharvesting bias [47], staying longer in patches than optimal (Figure 1d, dot-dash line).

To quantify subjects’ choice variability, we calculated each subject’s standard deviation of leaving behaviour, *SD*_*leave*_, for each patch type and environment (Figure 2a, Supplementary Figure 1a). We found that *SD*_*leave*_ in all tasks was relatively high, typically between 1 to 5 seconds, equating to about 20 to 50% of the average leaving time for that patch type in that environment. Thus subjects showed variable leaving behaviour in each combination of patch type and environment, consistent with them making stochastic leaving choices in identical contexts.

**Figure 2:**
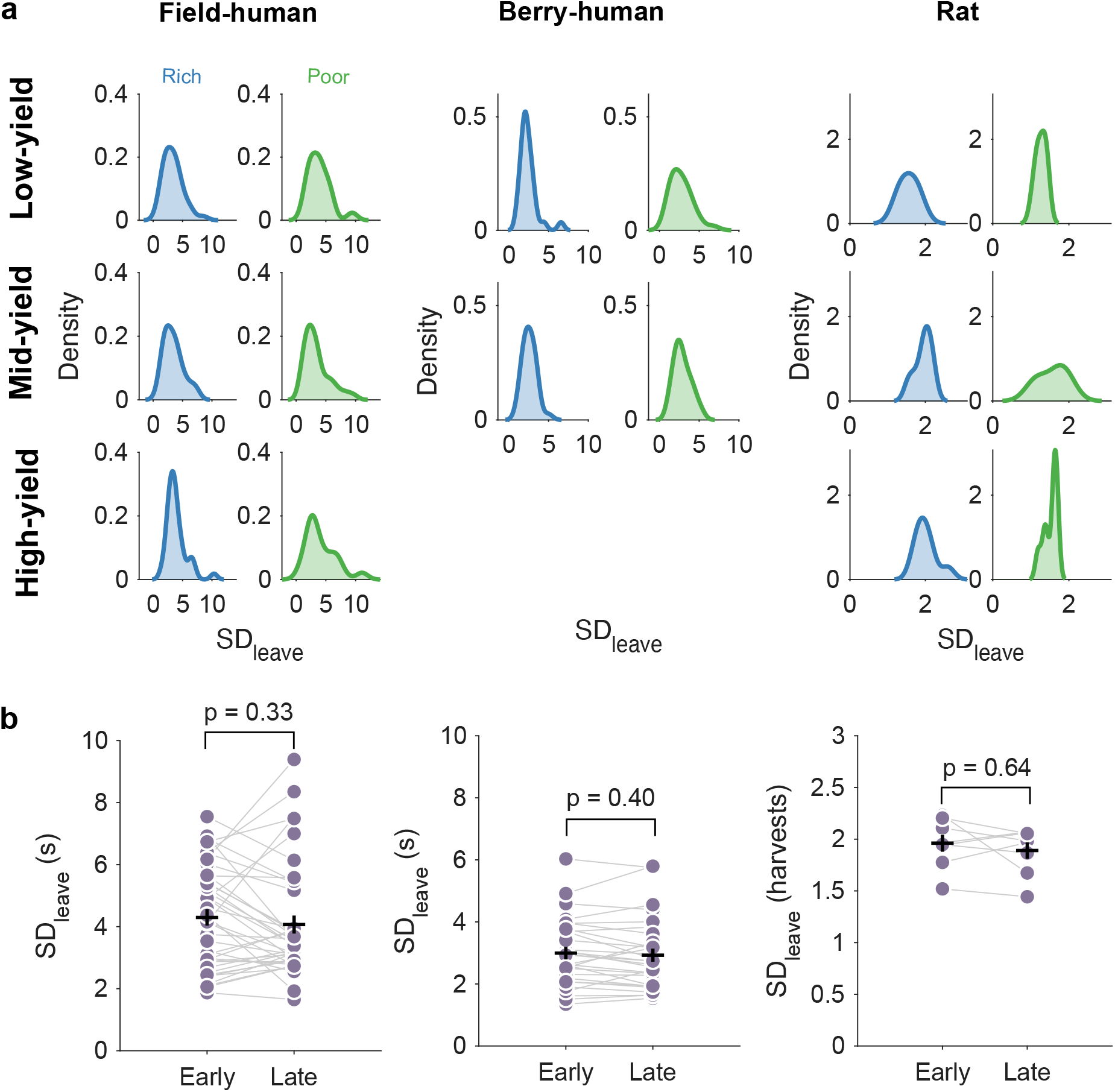
High variability in leaving behaviour across tasks. **a**: Distributions of subjects’ standard deviation in leaving behaviour, *SD*_*leave*_, for each patch and environment. Distributions are kernel density estimates. **b**: Subjects’ average *SD*_*leave*_ for early and late patch visits within each environment. Dots show *SD*_*leave*_ for each subject in one environment, grey lines show paired data points for each subject (field-human: n = 39, berry-human: n = 29, rat: n = 8). Early versus late trials are defined as the first versus second half of the time in an environment. Black horizontal line shows the mean *SD*_*leave*_, error bars show one SEM. p-values from Student’s t-test or Wilcoxon signed rank test (exact) on *SD*_*leave*_ between early and late patch visits.

We then asked if their variability was consistent over time, as would be expected from a fixed stochastic decision rule. We found that *SD*_*leave*_ was consistent across early and late visits to the same patch type within an environment (Figure 2b). We found no significant difference in *SD*_*leave*_ for early versus late patch visits in the field-human (*t*(38) = 0.98, *p* = 0.33, 95% CI = [-0.24, 0.70], Cohen’s *d* = 0.13) berry-human (*t*(28) = 0.85, *p* = 0.40, CI = [-0.09, 0.23], Cohen’s *d* = -0.06) and rat task (*W* (7) = 22, *p* = 0.64, CI = [-0.12, 0.15], *d* = 0.20). This absence of change in variability was supported by Bayes factors for the null hypothesis (*BF*_01_) of 3.71 for the field-human task, 3.64 for the berry-human task, and 2.55 for the rat task, meaning the data were 3.71, 3.64 and 2.55 times more likely under the null hypothesis of no effect of early versus late patch visits on *SD*_*leave*_. Subjects’ leaving time variability was thus consistent over time, suggesting a fixed stochastic contribution to the stay-or-leave decision itself.

### Stochastic choice predicts variability should be largely independent of reward

To test our hypothesis that a stochastic decision rule contributes to variability in patch leaving, we asked what such stochastic choice predicts for the patterns of variability that foragers should show across patch and environment types in our three tasks. Motivated by stochastic action selection models in reinforcement learning [28], we studied stochastic stay-or-leave models of the general form

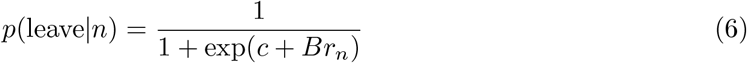

that give the forager’s probability of leaving a patch in time step *n* given the current reward obtained in the patch, *r*_*n*_. Using this general form allowed us to consider both reward-independent (*c*) and reward-dependent (*B*) contributions to stochastic choice: a reward-independent bias (*c*) modelling decision bias, lapses, and other non-reward contributions, with more negative values favouring leaving; and the forager’s relative weighting (*B*) of its own information about reward, which can be 0 or any positive number – the model thus gives an increasing probability of leaving as *r*_*n*_ decreases. Analysis of the model showed that, as expected, increasing either *c* or *B* increases the forager’s expected leaving time and that the model can recover the optimal leaving times predicted by the marginal value theorem (Supplementary Figure 2).

At first glance this recovery may seem odd as the marginal value theorem and its variant models (e.g. [16]) make a stay-or-leave decision by comparing the immediate patch reward *r*_*n*_ and the average reward rate *ρ* in the environment and our general form (Eq 6) does not. However, we initially assumed only that one or both of the bias (*c*) and weighting (*B*) parameters are influenced by the average reward rate, and so must differ between environments. For example, in the richer of two environments, we assumed the forager will have lower reward-independent bias (lower *c*), lower weighting of reward information (lower *B*), or both (Supplementary Figure 2). Below we will show that the data we study here are consistent with the average reward rate *ρ* defining the weighting *B* of patch reward information. But we first take advantage of the general form to make predictions for the variability of foragers’ stay-or-leave decisions without needing to commit to a specific model for the average reward rate.

In Figure 3 we plot these predictions for a forager’s variability in our three tasks. For the two human tasks with an exponential decay of reward our model predicts that a forager’s variability increases monotonically with the weighting of reward information (*B*) but plateaus at higher weightings (Figure 3a, c). By contrast, for the rat task with a linear decay of reward our model predicts that a forager’s variability is a non-monotonic function of the weighting of reward information (*B*), peaking at intermediate weightings (Figure 3e). Stochastic choice thus predicts that a forager’s leaving variability depends on how the resource decays when harvested.

**Figure 3:**
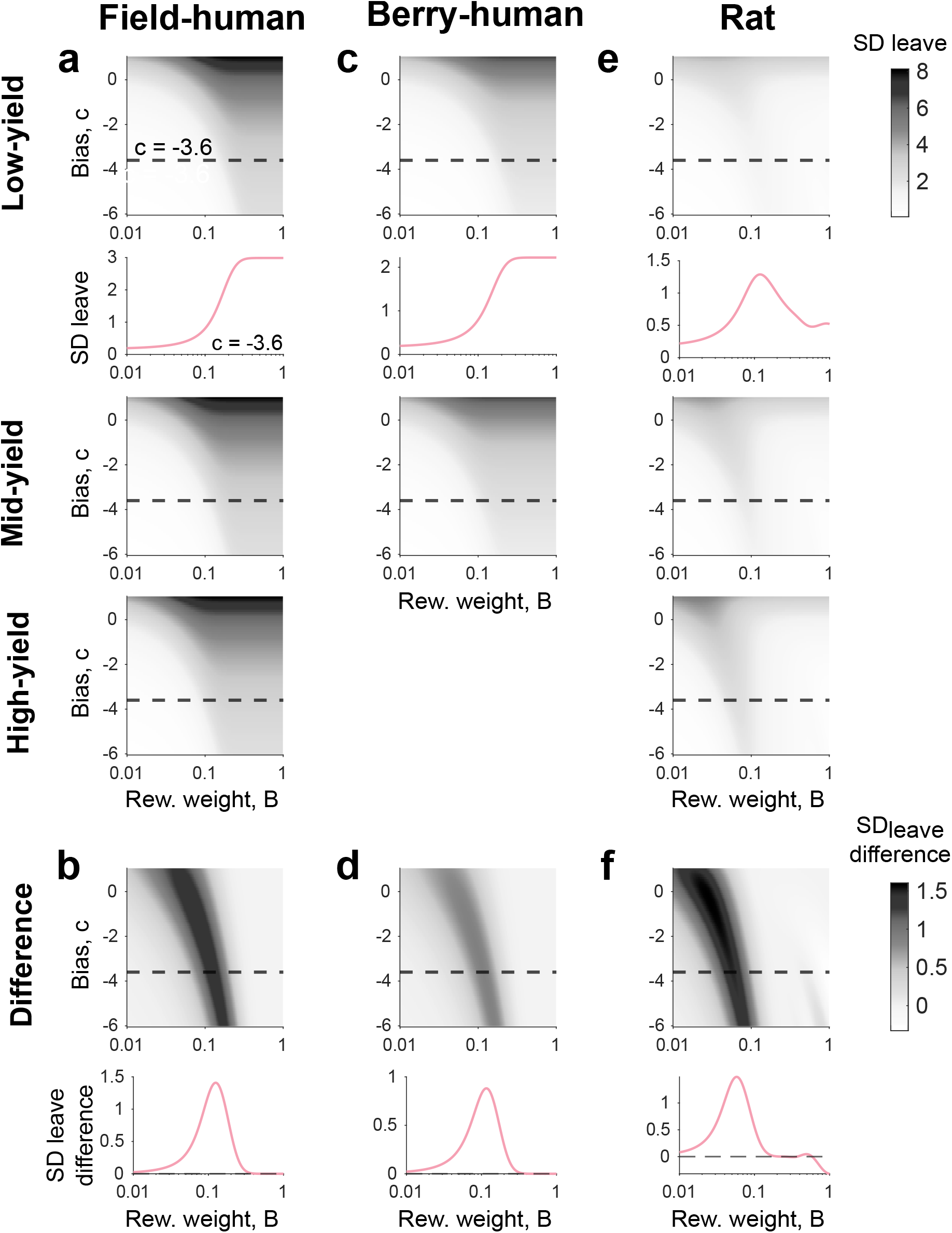
Stochastic choice predicts leaving variability depends on how reward decays. **a**: Heatmap of predicted *SD*_*leave*_ for different patch types in the field-human task, as a function of reward weighting (*B*, x-axis in log-scale) and reward-independent bias (*c*). Dashed line is the value of *c* for the inset plot of *SD*_*leave*_. Top right colourbar applies to panels a, c and e. **b**: Heatmap of predicted difference in *SD*_*leave*_ between high and low yield patches, for field-human task. Dashed line and panel inset as per panel a. Bottom right colourbar applies to panels b, d and f. **c-d**: As per a-b, for berry-human task. Since there are only two patch types, difference is between mid yield patch minus low-yield patch. **e-f** : As per a-b, for rat task.

The model further predicts that, in all tasks, there is a narrow range of reward-weighting (*B*) for which there is more choice variability in high yield than low yield patches, but that otherwise the forager’s variability does not differ between types of patch (Figure 3b, d, f). These predictions are the same at any chosen value for the reward-independent bias (*c*). Thus, the stochastic choice model predicts that, for a wide range of both the environment’s and forager’s parameters, there is no effect of reward on the variability of a forager’s choice.

### Real foragers’ variability can be independent of patch and environment

To test whether real foragers matched these predictions we fitted variants of our stochastic model to individual subjects’ behaviour in our three tasks. As our general model (Eq 6) assumes the average reward rate, which varied between environments in our tasks, contributes to either the reward-independent bias (*c*) or the weighting of reward information (*B*) or both, we fitted three model variants that allowed *c, B*, or both to vary between environments for each subject (Methods).

Fitting these models to subjects’ trial-by-trial leaving times provided good replications of the mean leaving time of the group and of each individual subject (Supplementary Figure 3a-f). Further, the model did not systematically over or under-estimate leaving times across the two environments (Supplementary Figure 3g-i). These models also matched subjects’ individual *SD*_*leave*_ (Supplementary Figure 4). We thus turned to the models’ subject-specific predictions for how variability would change across patch types and environments.

For the human tasks with exponentially-decaying rewards, fitting all three model variants found values of *c* and *B* that predicted subjects’ variability would be the same across all types of patch (Figure 4a and b shows results of the *‘vary B, fix c’* model; Supplementary Figure 5a-b, d-e shows results for the other two model variants). The models made different predictions for how the environment influences variability depending on subjects’ *c* or *B*. The two model variants that varied reward-weighting *B* between environments predicted that variability would be the same between environments (Figure 4a, b; Supplementary Figure 5d, e). The third variant, that varied only reward-independent bias, *c*, predicted higher variability in poor environments (Supplementary Figure 5a and b).

**Figure 4:**
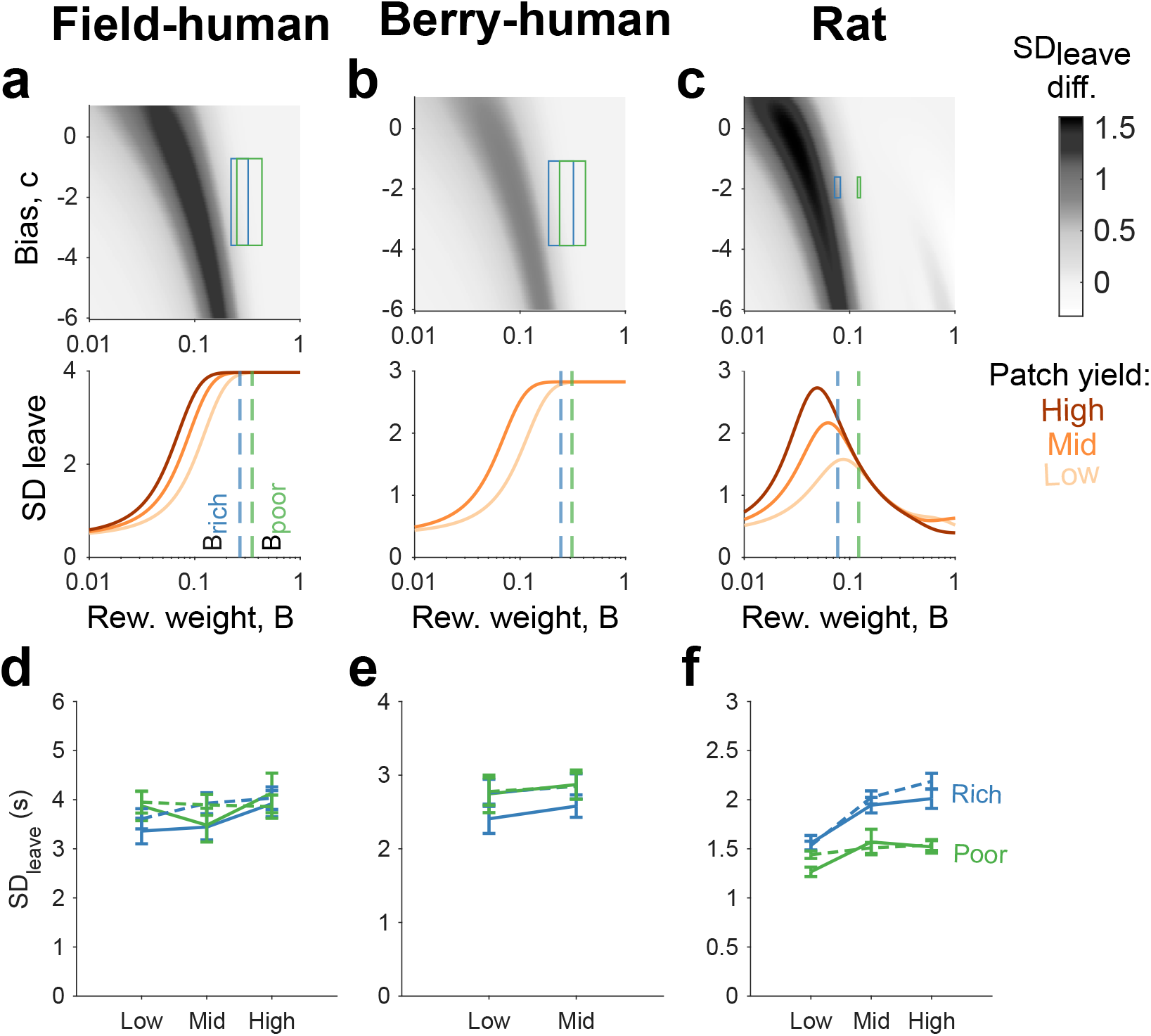
Subjects’ variability is consistent with model-predicted variability. Here we plot results for the *‘vary B, fix c’* model variant; Supplementary Figure 5 shows similar results for the other model variants. **a** Top panel shows the heatmap of predicted difference in *SD*_*leave*_, as per Figure 3a, overlaid with boxes showing 25th-75th percentile of subjects’ parameter fits for rich environment (blue) and poor environment (green). Bottom panel shows *SD*_*leave*_ as a function of *B* across different patch types, holding *c* fixed at its median fitted value. Dashed lines show the median fit *B* in rich (blue) and poor (green) environments. Colourbar applies to all heatmaps. **b**: As per panel a, for the berry-human task. **c**: As per panel a, for the rat task. **d** Group means of average *SD*_*leave*_ for the field-human task for subjects (solid line) (n = 39) and model simulations (dashed line), for each patch type and environment. **e**: As per panel d, for the berry-human task (n = 29). **f** : As per panel d, for the rat task (n = 8).

We found these predictions that variability could be the same across patches or environments counter-intuitive: because foragers in these tasks stay longer in higher-yielding patches and in poorer environments (Figure 1d) this implies that foragers decouple their leaving variability from how long they stay in a patch. Thus all model variants predicted we should find no difference in foragers’ average variability for leaving different types of patch, and the model variants that varied *B* between environments predicted we should find no difference in foragers’ average variability across different environments (Figure 4a and b), even though their average leaving times differ between patch types and between environments.

Strikingly, as predicted, the group averages of subjects’ empirical and model-estimated *SD*_*leave*_ showed little apparent change between types of patch or between environments (field-human: Figure 4d, berry-human: Figure 4e). Statistical analysis of the subjects’ empirical *SD*_*leave*_ further confirmed they showed little variation between patch types or environments (Supplementary Table 2, Supplementary Note 2). Together, these results show that, as predicted by variants of our stochastic choice model, subjects’ variability was independent of the type of patch or environment they were in, despite strong modulation of their *average* leaving times by both the patch and environment (Figure 4d and e). This also suggested that varying reward weighting between environments provided a better account of foragers’ behaviour than varying reward-independent bias, which we come back to below.

### Linearly-decaying rewards cause greater variability in rich environments

The model made strikingly different variability predictions for the rat task, which had linearly-decaying rewards. The values of *c* and *B* found from model fitting again well-replicated individual means (Supplementary Figure 3) and variations in leaving times (Supplementary Figure 4). All model variants now predicted that the rats’ variability would be sensitive to available patch reward, with low yield patches having a lower variability than other patch types (Figure 4c, Supplementary Figure 5). The two model variants that varied reward-weighting *B* between environments also predicted that the rats would have higher variability in rich than in poor environments (Figure 4c). We termed this an inverted environment effect, as the model predicts that the shorter average leaving times in the rich environment should correspond to *greater* variability of leaving times. The third model variant, that varied only reward-independent bias *c* between environments, did not predict this inverted environment effect (Supplementary Figure 5).

Nonetheless, despite these strikingly different predictions of the reward-dependence of variability compared to tasks with exponentially-decaying reward, we found the group averages of subjects’ empirical and model-estimated *SD*_*leave*_ matched them, with low-yield patches having the lowest variability and rich environments having a higher variability of leaving times than poor environments (Figure 4f). Statistical analysis of the rats’ empirical *SD*_*leave*_ confirmed both results, including their greater variability of leaving times in the rich environment (Supplementary Table 2, Supplementary Note 2), again suggesting that varying reward weighting rather than bias between environments provided a better account of foragers’ behaviour, to which we turn next.

### The average reward rate controls the weighting of reward information

The above established that stochastic choice predicts the patterns of foragers’ leaving variability across patch types and environments, and does so across tasks and species. Each variant of the model tested so far embodied a different hypothesis about how the environment’s average reward rate *ρ* affected that choice: through the reward-independent bias (*c*), the reward weighting (*B*), or both. To distinguish these hypotheses we compared the quality of their fits to the data. Model selection metrics showed the best-fitting of these models at the group (Figure 5a) and individual (Supplementary Figure 6a) level was consistently *‘vary B, fix c’* across the three tasks, supporting the hypothesis that the average reward rate affects stochastic choice by controlling the weighting of reward information.

**Figure 5:**
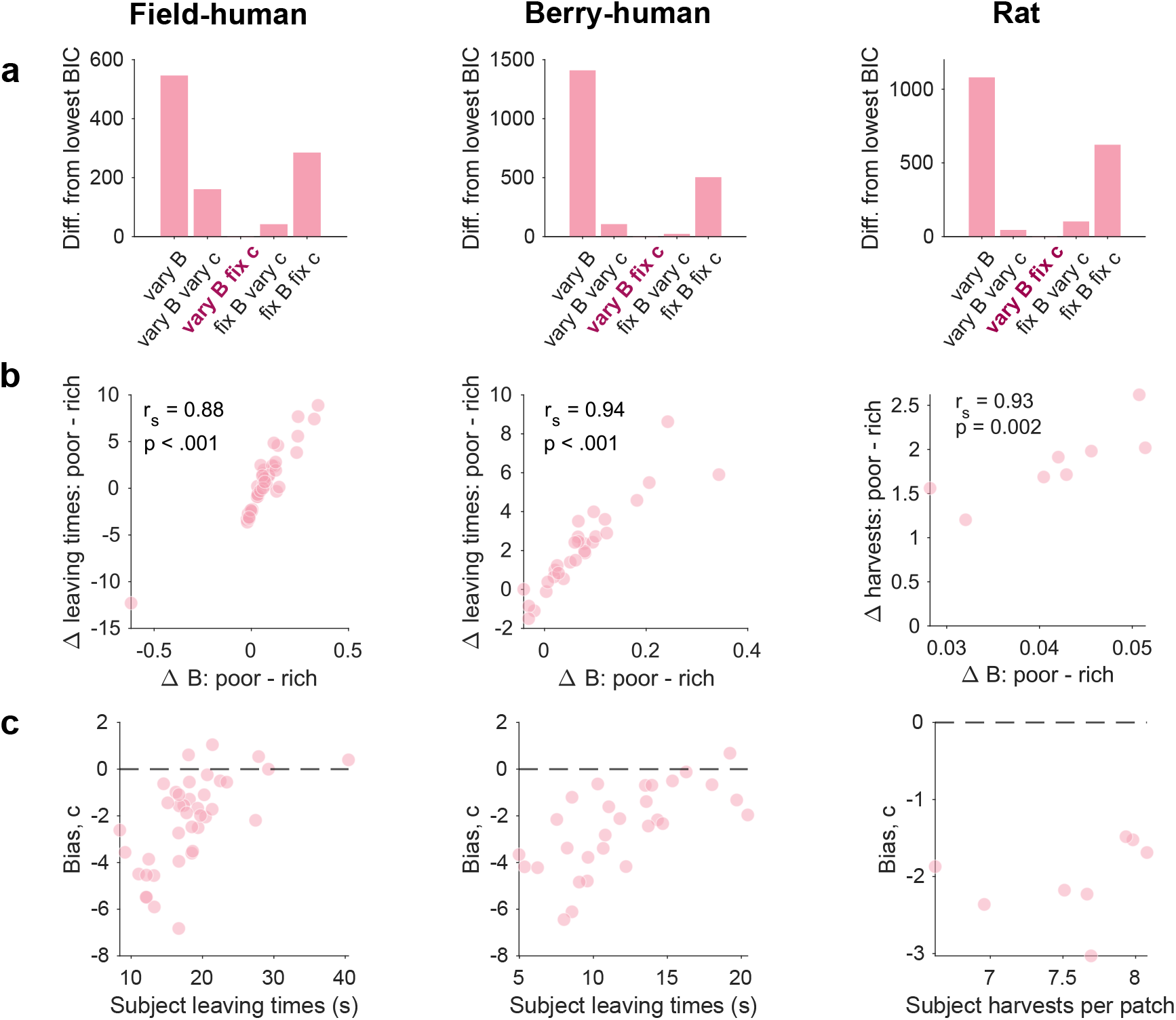
Subjects’ leaving behaviour is best explained by *‘vary B, fix c’* model: environment-dependent reward weighting plus a consistent bias. **a**: Model comparison using Bayesian Information Criterion (BIC). We plot for each model the difference between its BIC and the BIC of the highlighted winning model (the model with the lowest BIC, ‘vary B fix c’). **b**: Correlation of difference in *B* fits between environments (Δ*B*) and difference in mean leaving times between environments (Δ leaving times). *r*_*s*_: Spearman’s correlation coefficient (field-human: n = 39, berry-human: n = 29, rat: n = 8). **c**: Scatterplot of subject leaving times/number of harvests with *c* fits for each subject. Black dashed line at *c* = 0 shows where the reward-independent bias switches from leaving to staying.

We further explored evidence for this hypothesis by considering two additional control models. The first (*‘fix B, fix c’*) tested the hypothesis that the average reward rate *ρ* does not affect choice, so had the same reward weighting *B* and reward-independent bias *c* between environments. A substantial minority of individual subjects were best-fit by this model (Supplementary Figure 6a) rather than *‘vary B, fix c’*. Inspection of the data showed that which of these two models best-fit an individual simply depended on whether or not they showed an effect of the environment on their behaviour: our model predicts that subjects showing little effect of the environment on their mean leaving times would have the same weighting of reward information in those environments – and so would be better fit by a model with fixed reward weighting *B*. We indeed found that the effect of the environment on a subject’s mean leaving time strongly and positively correlated with the difference in fitted *B* between environments in the *‘vary B, fix c’* model (Figure 5b). These data thus suggest that, while subjects do use the average reward rate to set their reward weighting, there is variation in how they do so, which we take up below.

The second control model (*‘vary B’*) tested the hypothesis that subjects did not have a reward-independent bias (*c* = 0), such that their behaviour could be explained entirely by their reward weighting *B* changing between environments. This model was consistently the worst-fitting of all tested (Figure 5a), implying that most subjects had a consistent bias towards staying or leaving, independent of reward. That consistent bias was towards leaving: the fitted values of *c* for the *‘vary B, fix c’* were negative for almost all subjects across the three tasks (Figure 5c).

To confirm each of these model inferences were robust, we verified that *‘vary B, fix c’* and both control models were uniquely identifiable among the models we tested (Supplementary Figure 6b) and that the *‘vary B, fix c’* model’s parameter values were recoverable with high accuracy (Supplementary Figure 6c), given the quantity of data used to fit the models and the found parameter ranges for each task. Together, these results support two hypotheses: first, that real foragers make stochastic leaving choices driven by both reward-independent bias and a weighting of reward information; and, second, that foragers vary in the degree to which the environment’s average reward rate controls the weighting of reward information.

### Foraging choice can be explained by a subject-specific internal function for reward weighting

This second hypothesis implies that foragers have an internal function for how they translate average reward rate *ρ* in the environment into their weighting *B* of reward information in a patch. To test this, we asked whether a model that explicitly derives the reward weighting from the average reward rate could account for subjects’ behaviour in our tasks. We fitted four different internal functions to subject data (*B* M1 - M4, described fully in Methods) that made leaving choices as above but derived a subject’s *B* as a function of their experienced average reward rate in the environment, *ρ*. Given that *B* is higher in poor environments (Supplementary Figure 6d), we formulated our internal functions so that lower average reward rates led to higher values of *B*. The goal was to test our internal function hypothesis by asking if there existed an internal function of the form *B* = *f* (*ρ*) that could provide a good account of subjects’ behaviour; we thus used model selection to identify good models for further analysis, not to support a specific hypothesis for this internal function based on the ‘best’ model.

We found that internal functions of the general form *B* = *λρ*^*γ*^, which included both M1 and M4, well explained subject behaviour (Supplementary Figure 7a). The berry-human and rat subjects were best fit by the general form of this model, M4, where *γ* could vary freely across subjects (Figure 6a); the field-human dataset was best fit by the simpler version of this model, M1, where *γ* was fixed (*γ* = −1) across subjects rather than allowed to vary (Figure 6a). Since M1 and M4 have the same general form, we interpret these differences in winning models across the datasets as small variations within the same model, rather than qualitatively different accounts of the internal function mapping average reward and *B*. Thus we continued our analysis using the general form of the model, M4. This general model almost perfectly replicated group-level (Figure 6b,c) and individual (Supplementary Figure 7b,c) leaving times and their variation across all tasks, including the decoupling of leaving variation from reward in the human tasks and the inverted environment effect on leaving variation in the rat task (Figure 6c).

**Figure 6:**
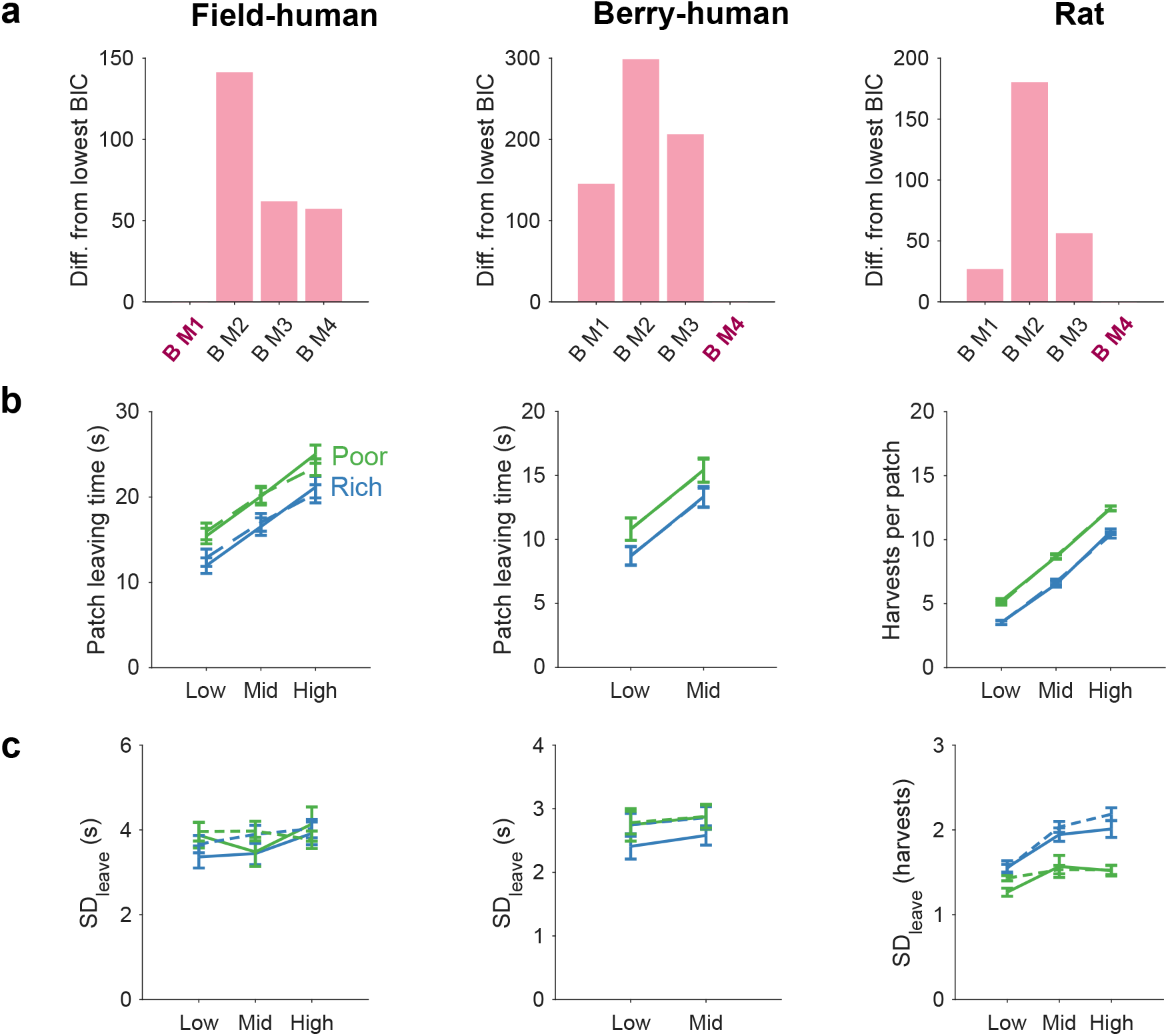
Reward weighting is a function of the environment’s average reward rate. **a**: Model comparison using Bayesian Information Criterion (BIC). Models M1 to M4 differed in the function linking the average reward rate to forager’s weighting of reward information (*B*) in their stochastic choice to leave. We plot for each model the difference between its BIC and the BIC of the highlighted winning model (the model with the lowest BIC). **b**: Group means of leaving times/number of harvests for the subjects (solid line) (field-human: n = 39, berry-human: n = 29, rat: n = 8) compared to simulations of M4, (dashed line) for each patch type and environment. **c**: Group means of *SD*_*leave*_ for subjects (solid line) compared to simulations of M4 (dashed line), for each patch type and environment.

These results show a forager’s stochastic leaving choice could be defined by a single set of behavioural parameters that were fixed across environments: the forager’s reward-independent leaving bias (*c*), and two parameters (*λ* and *γ*) that define their internal function for translating their experienced average reward rate into the weighting of patch reward. Our model fitting showed that both *γ* and *λ* parameters vary between subjects and correlate with the difference in leaving behaviour between rich and poor environments (Supplementary Figure 7d,e), consistent with our hypothesis that the internal function varies between subjects.

Whilst the subjects in our tasks were well-practised, and thus we assumed in the above model a stable estimate of the average reward rate, typically the environment’s statistics are not initially known to the forager and must be learned through experience. To check our internal function hypothesis was robust to learning, we extended the general model (M4) to learn *ρ* per time-step (Methods; Supplementary Figure 8) and found it still accurately reproduced forager’s leaving time and variability. Thus, a stochastic choice model that updates its weighting of reward over time could still account for variability in patch leaving across different task structures and species.

## Discussion

Patch foraging is a fundamental decision problem across species, requiring a choice between staying to harvest known remaining resources or leaving to find another patch of unknown quality [8, 19, 9]. These ‘stay-or-leave’ decisions are traditionally compared to the marginal value theorem [10], which assumes deterministic leaving decisions. Here we showed that, across tasks and species, foragers were highly variable in their decision of when to leave and that this variability was consistent with them making stochastic leaving decisions.

A striking prediction of our stochastic choice model is that the variability of patch leaving times is constant across different patch values or richness of the environment for a wide range of its parameters. Within those ranges, foragers are predicted to show a decoupling of leaving variability from how long they stayed in the patch. This is in stark contrast to reaction-time and evidence-accumulation decision tasks where the variability in decision time is directly proportional to its mean [48]. We revealed here nonetheless that human foragers in two tasks unexpectedly showed exactly this predicted uncoupling, with no change in their variability of leaving times across patch types and environments, despite changes in the value of reward and of their mean leaving times.

Notably, in the third task, with rats experiencing linear decay of reward in each patch, subjects’ variation in leaving times *did* depend on both the value of the patch and the richness of the environment. Comparing only these empirical differences in variability between human and rat tasks would have supported an inference of different decision processes for leaving between tasks and species. But our stochastic choice model had predicted exactly the pattern of leaving time variability shown by the rats across patch types and environments, evidence that the same stochastic choice model underlies variable leaving decisions across species.

The data here supported the further hypothesis that this common model of stochastic choice combined both an environment-dependent weighting of reward and a constant reward-independent bias towards leaving. These results align with existing models of patch foraging, where a decision bias term improved the model fit for human [16] and rat [15, 49] patch leaving behaviour. However, these latter results found that rats had a different bias between environments. By including both reward weighting and reward-independent bias together, our results alternatively suggest that it is changes in the weighting of patch reward, *B*, that drive environment differences in patch leaving choices, while reward-independent biases remain consistent across environments. This bias could be modelling one or more factors that do not change between environments, including motor, sensory or learning noise [50, 51], inattention [52], and risk aversion [16, 23].

We showed that the change in reward weighting between environments is consistent with foragers’ having an internal function for converting an environment’s average reward rate into that weighting. Miscalibration of this internal function could account for overharvesting. Foragers of a wide-range of species typically overharvest when foraging [47], including in the tasks we studied. There are several computational accounts for overharvesting, including delay discounting, where individuals prefer immediate over delayed reward [23], sensitivity to sunk costs in the current patch [49] or risk aversion, where the marginal value of a new patch is lower than the current one [16]. These models propose that foragers generally try and follow a deterministic, optimal rule, but are influenced by fixed decision biases that enforce staying in patches longer. Our miscalibration hypothesis instead predicts that a forager’s overharvesting arises from its internal function converting average reward rate into too high a weighting of patch reward. This aligns with findings that overharvesting may stem from inaccurately learning the structure of the environment [53] or the average reward rate [26]. However, our results suggest that, even with perfect information about rewards in the patch and environment, a forager could still display overharvesting as a result of a miscalibration in their internal function for translating reward into stochastic choice.

An interesting question is why foragers would make a stochastic choice of when to leave a patch. The stochastic choice model we use here was inspired by the softmax action selection model in reinforcement learning [28], where the inverse temperature parameter (*B*) is widely interpreted as controlling the exploration-exploitation trade-off in action selection. Discussion of an exploration-exploitation trade-off is also common in the foraging literature, but the two uses typically refer to different concepts: in reinforcement learning, the explore-exploit trade-off refers to how much the probability of selecting an action is weighted by its predicted value, with high weightings leading agents to exploit actions with the currently known highest value and low weightings leading agents to explore other, potentially more valuable, actions [28, 3, 30]. In foraging, the exploit-explore trade-off refers to whether the forager should stay and exploit the resources in the current patch or leave and explore other patch options, with no reference to whether the decision is stochastic or variably weights reward information [54, 19, 55, 56, 20]. Our model asks if foragers combine the two concepts: if they control the weight of reward information when they make a stochastic decision to leave; our results suggest they do. Stochastic choice based on different weightings of reward might be suboptimal for patch foraging as it deviates from the deterministic choice of the marginal value theorem; but foragers’ use of stochastic choice could reflect a general mechanism of flexible action selection that animals apply across different types of decision making [57].

### Limitations

While we showed subjects’ variability is consistent with an internal function translating average reward into reward weighting *B*, we have not yet rigorously assessed the form of that function. We selected several simple functions linking average reward and *B*. However, there are many other mappings that could describe the relationship, which may have important theoretical or biological significance. Future work is required to characterise this mapping in more detail, identifying the best fitting function and what this means for the neural mechanisms of stochastic choice.

Both the human tasks had exponentially decaying reward and thus our model made the same, validated prediction: the decoupling of decision variability from patch type and environment. However, by deriving predictions for the model across a range of reward-weighting parameter *B*, we also showed that there are regimes of *B* where variability *is* coupled with patch type and environment. Specifically, our model predicts that lower values of *B* would lead to greater variability in poor compared to rich environments, and greater variability as patch yield increases (Figure 4). We were unable to test this further prediction with the datasets here. Thus, further rigorous testing of our model would test these predictions in a dataset characterised by lower levels of *B*. This would require developing a task in which participants left patches closer to the optimal leaving time.

Here we asked whether stochasticity in the decision process was sufficient to explain foragers’ variability in patch leaving. Variability may also arise from foragers’ uncertainty about parameters that feed in to the decision to leave: such uncertainty could lead to variable leaving times even when using a deterministic leaving rule. Uncertainty could arise from a number of sources, including estimating the patch or average reward rates [58, 20], inferring the structure of the environment [53], or from internal sensory uncertainty, such as estimating the passage of time [59]. Further, naturally-occurring stochasticity in the environment will produce uncertainty, requiring continual estimation of reward statistics [60]. In the tasks studied here, most sources of uncertainty would decrease over time and consequently so would variability driven by that uncertainty. Yet we found that variability remained consistently high over time, suggesting that uncertainty was not strongly influencing patch leaving decisions in these specific tasks. Future work is needed to explore how stochastic choice as predicted by our model and models of uncertainty interact during stay-or-leave decisions in foraging. Relatedly, future research could examine how variability in patch leaving is modulated by changes in psychological states other than uncertainty, such as boredom or fatigue in humans [31], and internal nutritional state in animals [61].

## Data Availability Statement

The source data used to produce the figures in this paper are available from GitHub (https://github.com/Humphries-Lab/Foraging-variability). Original source data for the berry-human task are available from the data repository of Contreras-Huerta et al [41], and the source data for the rat task are available from Figure 1 of [23].

## Code Availability Statement

The code to reproduce the analysis and figures for this paper are available from GitHub (https://github.com/Humphries-Lab/Foraging-variability).

## Author Contributions

Conceptualisation by E.V.S, M.A.J.A and M.D.H. Data analysis by E.V.S and M.D.H. E.V.S and M.D.H wrote the first draft of the manuscript; E.V.S, M.A.J.A and M.D.H edited the manuscript. Supervision by M.A.J.A. and M.D.H.

## Acknowledgements

This work was funded by an MRC IMPACT doctoral training scholarship (MR/N013913/1) to EVS; a Biotechnology and Biological Sciences Research Council David Phillips Fellowship (BB/R010668/2), a Jacobs Foundation Fellowship and a Wellcome Trust Discovery Award to MAJA; and a Biotechnology and Biological Sciences Research Council grant [BB/X013111/1] to MDH and MAJA. The funders had no role in study design, data collection and analysis, decision to publish or preparation of the manuscript.

