## Supplementary Information for "Stochastic choice drives variability in patch foraging decisions in humans and rats"

April 3, 2026

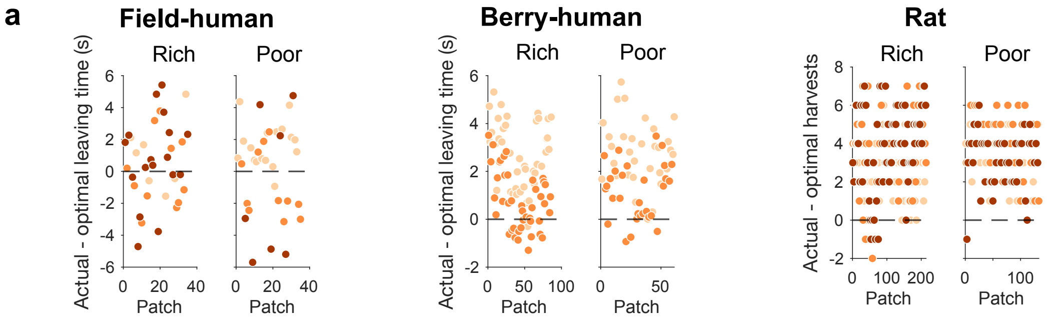


Supplementary Figure 1: **Variability in individual leaving behaviour and in different environments.** **a**: An example subject's deviation from optimal leaving behaviour in each environment, for each patch visited. Dots are coloured by patch type: red = high yield, orange = mid yield, beige = low yield. Black dashed line at $y = 0$ shows no deviation from optimal.

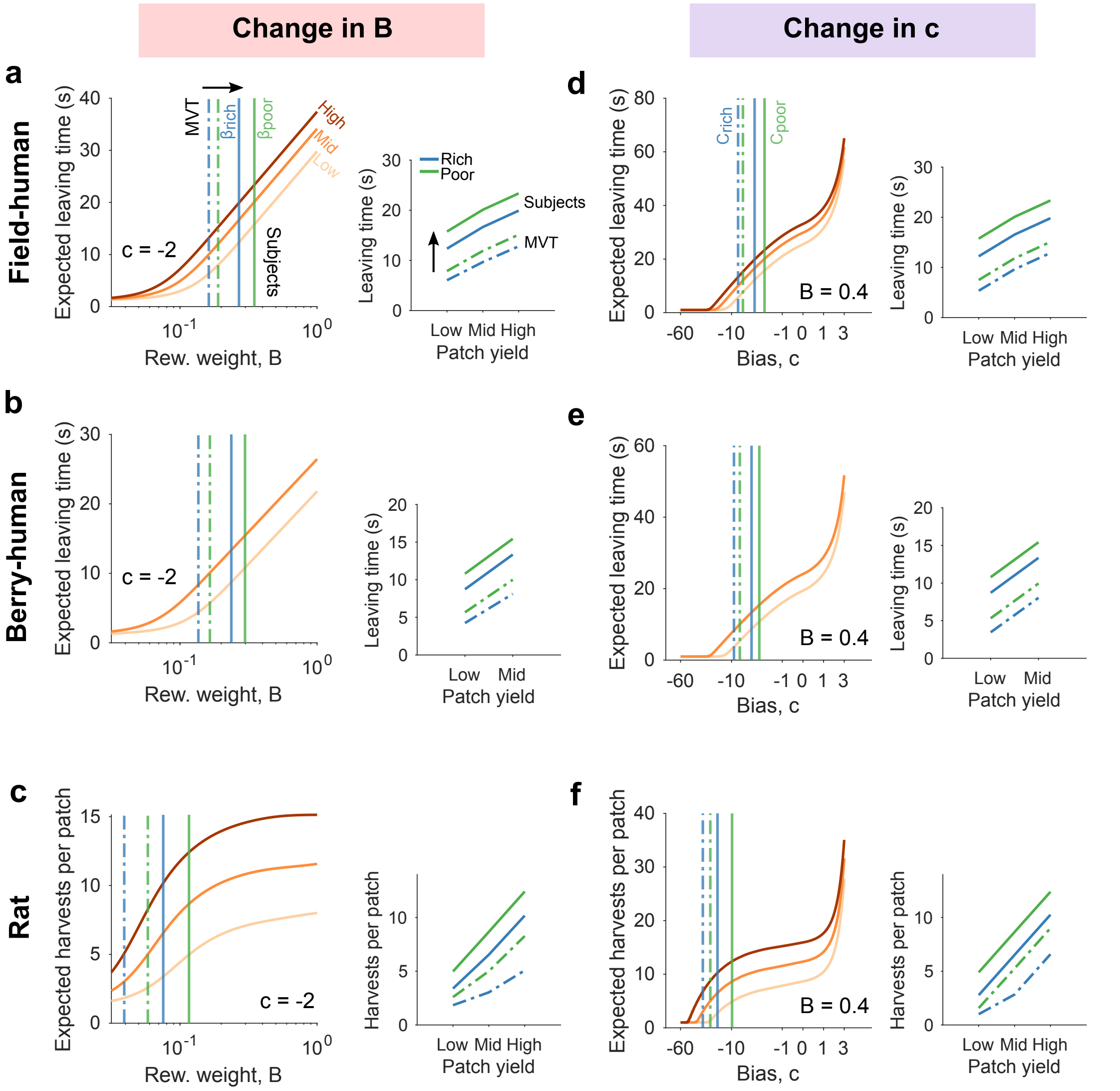


Supplementary Figure 2: **Model predictions for expected leaving behaviour a**: Expected leaving times of the model for the field-human task with exponential patch decay across a range of reward-dependent control $B$, with $c$ held fixed ($c$ = -2). X-axis ($B$) shown in log-scale. For all patch types, expected leaving times increase as $B$ increases (increased reward weighting). Dashed blue and green lines in the main panel show the values of $B$ in rich (blue) and poor (green) environments required to approximately produce optimal leaving times defined by MVT, for the mid-yield patch. Solid blue and green lines show the values of $B$ required to approximately produce mean subject leaving times for the mid-yield patch. Panel insets show expected leaving times of the model for MVT (dashed line) and subjects (solid line) at these parameter values, for each patch type and environment. **b-c**: as per panel a for the berry-human task (exponential decay) and rat task (linear decay) respectively. **d**: Expected leaving times of the model for the field-human task with exponential patch decay across a range of reward-independent bias $c$, with $B$ held fixed ($B = 0.4$). As per panel a, except blue and green lines show values of $c$ in rich and poor environments, required to produce MVT (dashed) and mean subject leaving times (solid). X-axis shown in signed log scale. Panel insets as per inset in panel a. **e-f**: as per panel d for the berry-human task (exponential decay) and rat task (linear decay) respectively.

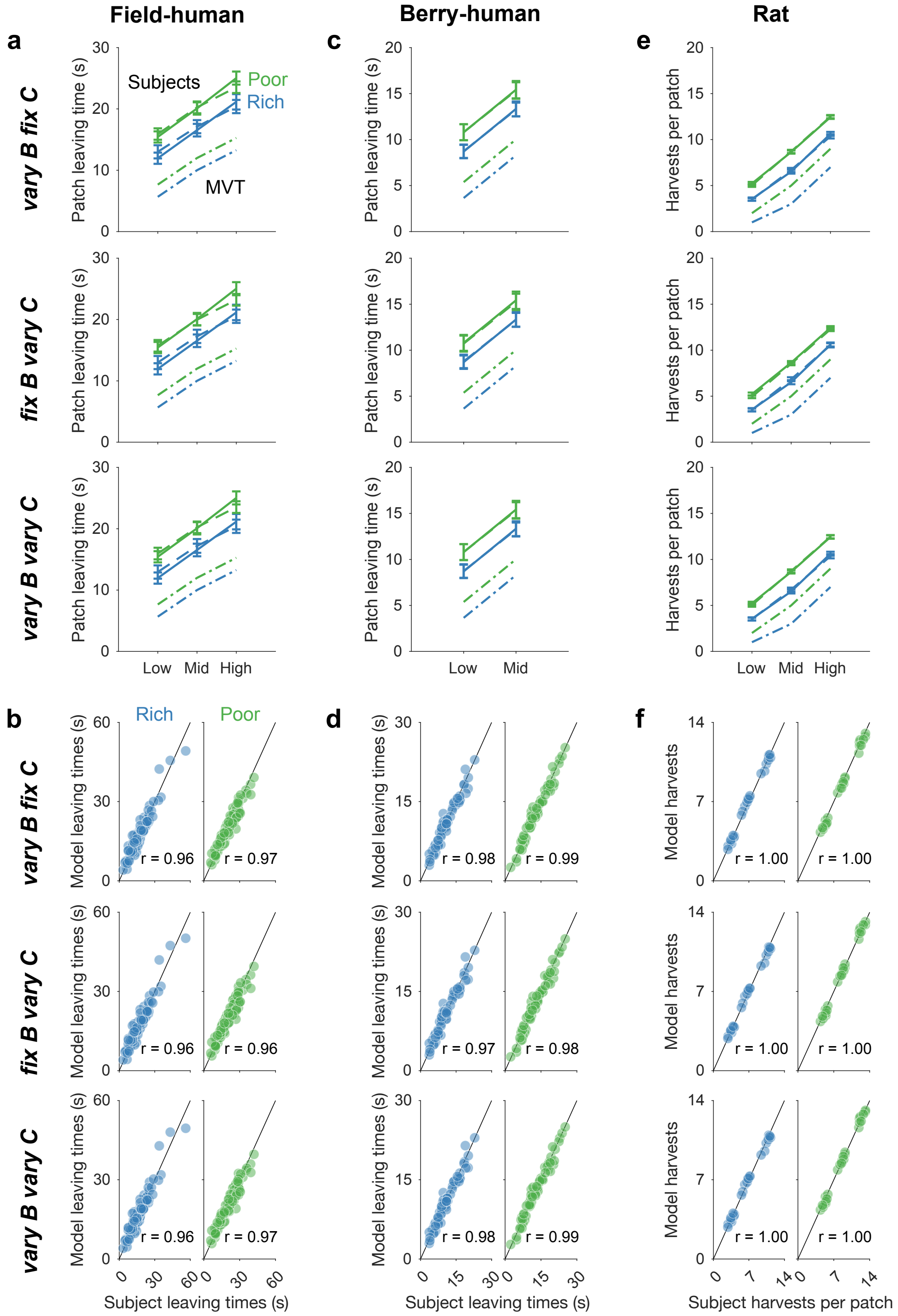


Supplementary Figure 3: **Subject's average leaving behaviour is captured by models of stochastic choice**. **a**: Simulated model leaving times for the field-human task, for each of three models (top: *'vary B, fix c'*, middle: *'fix B, vary c'*, bottom: *'vary B, vary c'*). Plots show group means of leaving times for the subjects (solid line) and for simulations of the fitted model (dashed line), compared to MVT predictions (dash-dot lines), for each patch type and environment. **b**: Correlation of subject and model leaving decisions, for each of the three models. Each dot is the mean leaving time/number of harvests for each subject and patch type, separated by environment. r: Pearson's correlation coefficient. **c, d**: as per panel a-b, for the berry-human task. **e, f**: as per panel a-b, for the rat task. Y-axis shows number of harvests per patch.

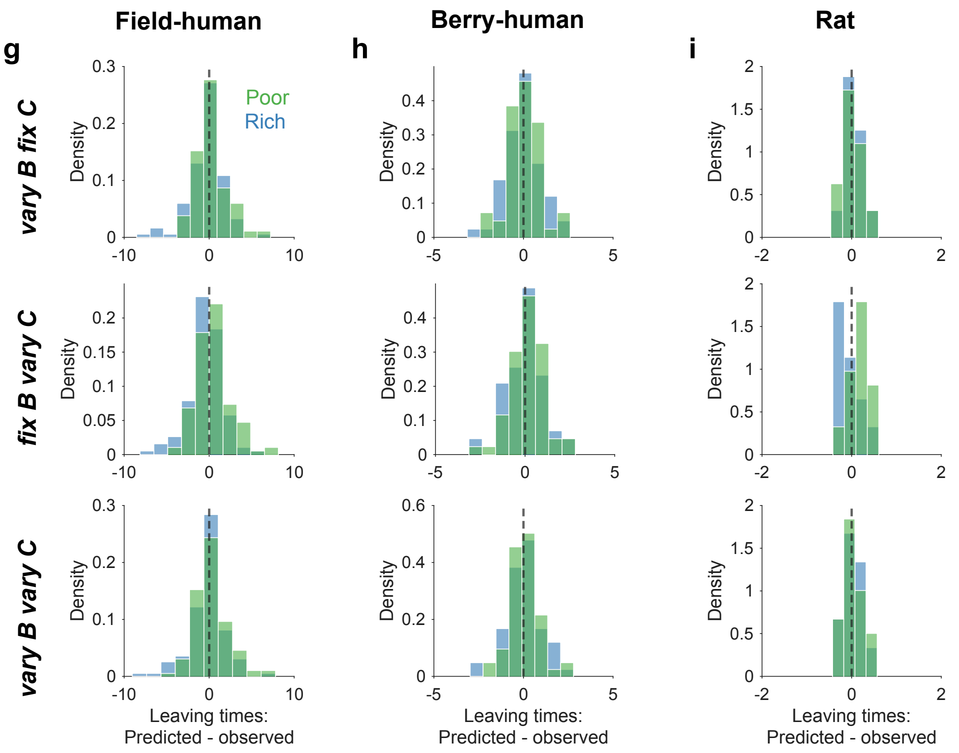


Supplementary Figure 3: **continued**. **g**: Distribution of differences in predicted versus observed leaving times for the field-human task, for each model (top: *'vary B, fix c'*, middle: *'fix B, vary c'*, bottom: *'vary B, vary c'*). Plots show probability density of differences for each environment (rich = blue, poor = green). **h**: As per panel g, for the berry-human task. **i**: As per panel h, for the rat task.

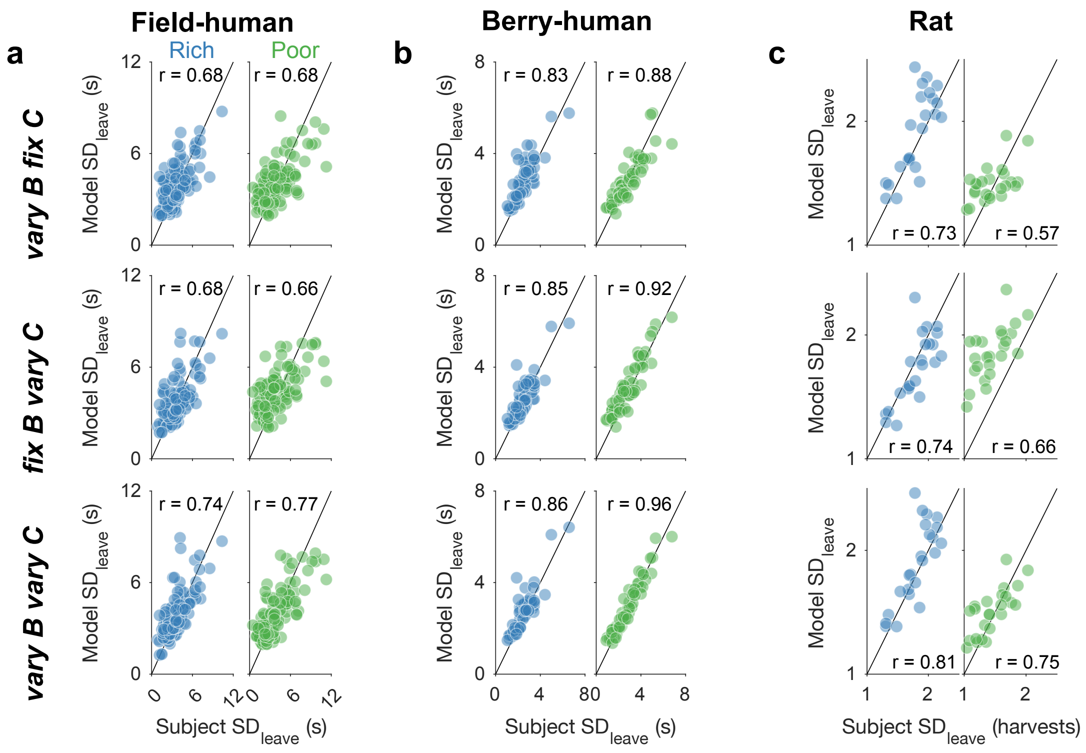


Supplementary Figure 4: **Subject's variability in leaving is captured by models of stochastic choice.** **a**: Correlation of average $SD_{leave}$ for each subject with average simulated $SD_{leave}$ for the subject. Each dot is the $SD_{leave}$ for each subject and patch type, separated by environment. r: Pearson's correlation coefficient. **b**: as per panel a, for the berry-human task. **c**: as per panel a, for the rat task. Y-axis shows SD for number of harvests per patch rather than leaving times in seconds.

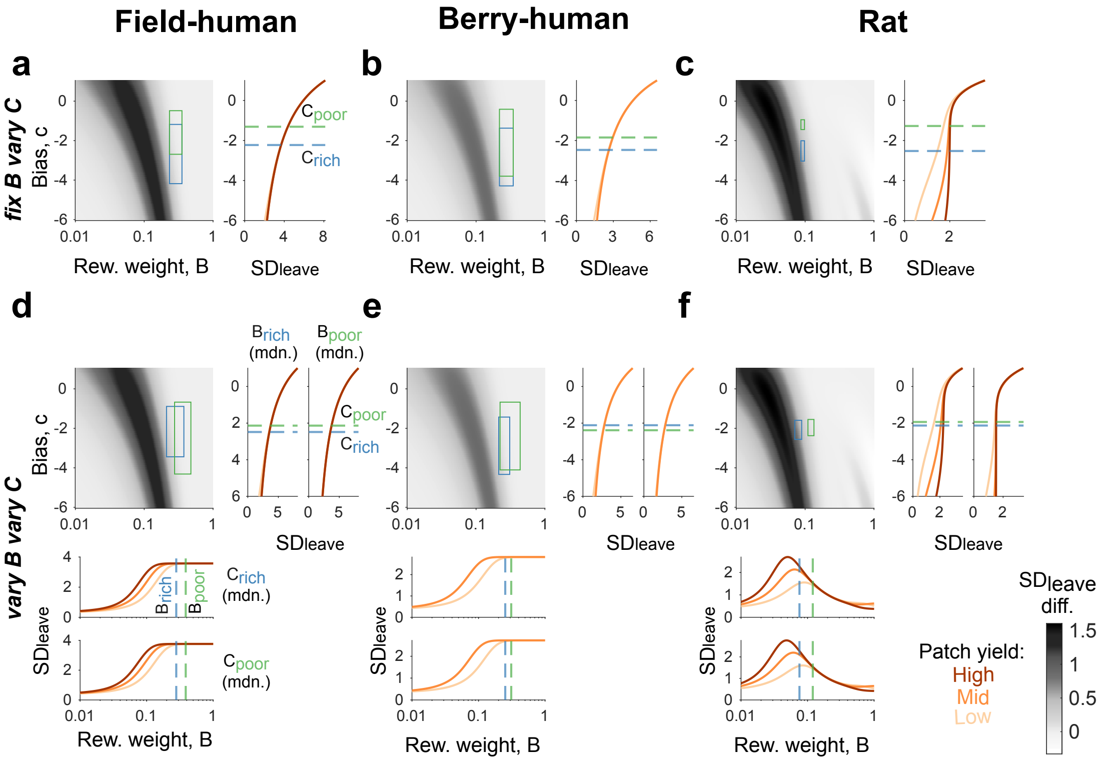


Supplementary Figure 5: **Model predicted variability for range of parameter fits, for *'fix B, vary c'* and *'vary B, vary c'* model.** **a**: Predictions of the *'fix B, vary c'* model for the field-human task. Left panel shows heatmap of predicted difference in $SD_{leave}$, overlaid with boxes showing 25th-75th percentile of subjects' parameter fits for rich environment (blue) and poor environment (green). Right panel shows $SD_{leave}$ as a function of $c$ across different patch types, holding $B$ fixed at the median fit. Note the curves for the three patch types exactly overlap, predicting identical $SD_{leave}$ across types. Lines show the median fit $c$ in rich and poor environments. Colourbar in bottom right applies to all heatmap panels a, b and c. **b**: As per panel a, for the berry-human task. **c**: As per panel c, for the rat task. **d**: As per panel a, for the *'vary B, vary c'* model. Right panel now shows separate predictions when fixing $B$ at the median subject fit for rich (left) and poor (right) environments. Bottom panel shows separate predictions when fixing $c$ at the median subject fit for rich (top) and poor (bottom) environments. Lines show median fits in each environment for the other parameter.

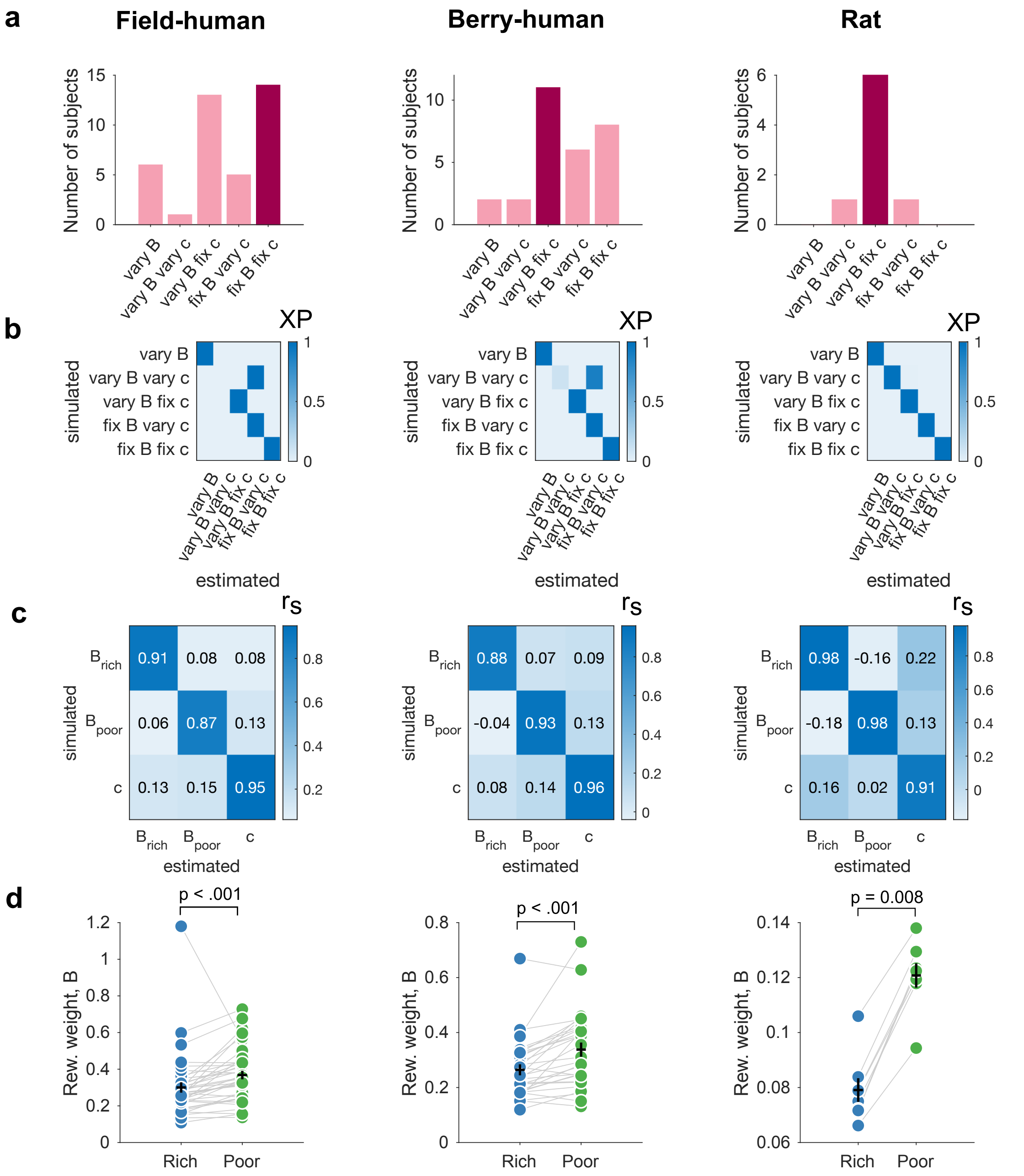


Supplementary Figure 6: **Additional model comparison metrics for stochastic models a**: Number of subjects best fit by each model, according to minimum BIC. Highlighted bar shows the model best fit for the highest number of subjects. **b**: Model identifiability matrix. $XP$: Exceedance probabilities indicating the confidence that the simulated data from each model is best fit by the same model compared to the other models tested. Deviations from the unit matrix for the *'vary B, vary c'* model indicates this model was better recovered by the simpler *'fix B, vary c'* model for the human tasks (n = 100 simulations per model). **c**: Parameter recovery matrix for winning model (*'vary B, fix c'*), showing correlation between simulated and recovered parameters. $r_s$: Spearman's correlation coefficient (n = 100 simulations). **d**: Estimated $B$ fits in rich and poor environments. Each dot represents a subject, grey lines show paired data points. Black horizontal line shows the mean $B$ per environment, error bars show 1 SEM. p-values from Student's t-test or Wilcoxon signed rank test (exact) on $B$ fits between rich and poor environments. Field-human: $W(38) = 709$, CI = [0.04, 0.09], $d = 0.77$, berry-human: $W(28) = 403$, CI = [0.03, 0.08], $d = 0.62$, rat: $W(7) = 36$, CI = [0.03, 0.05], $d$=5.54.

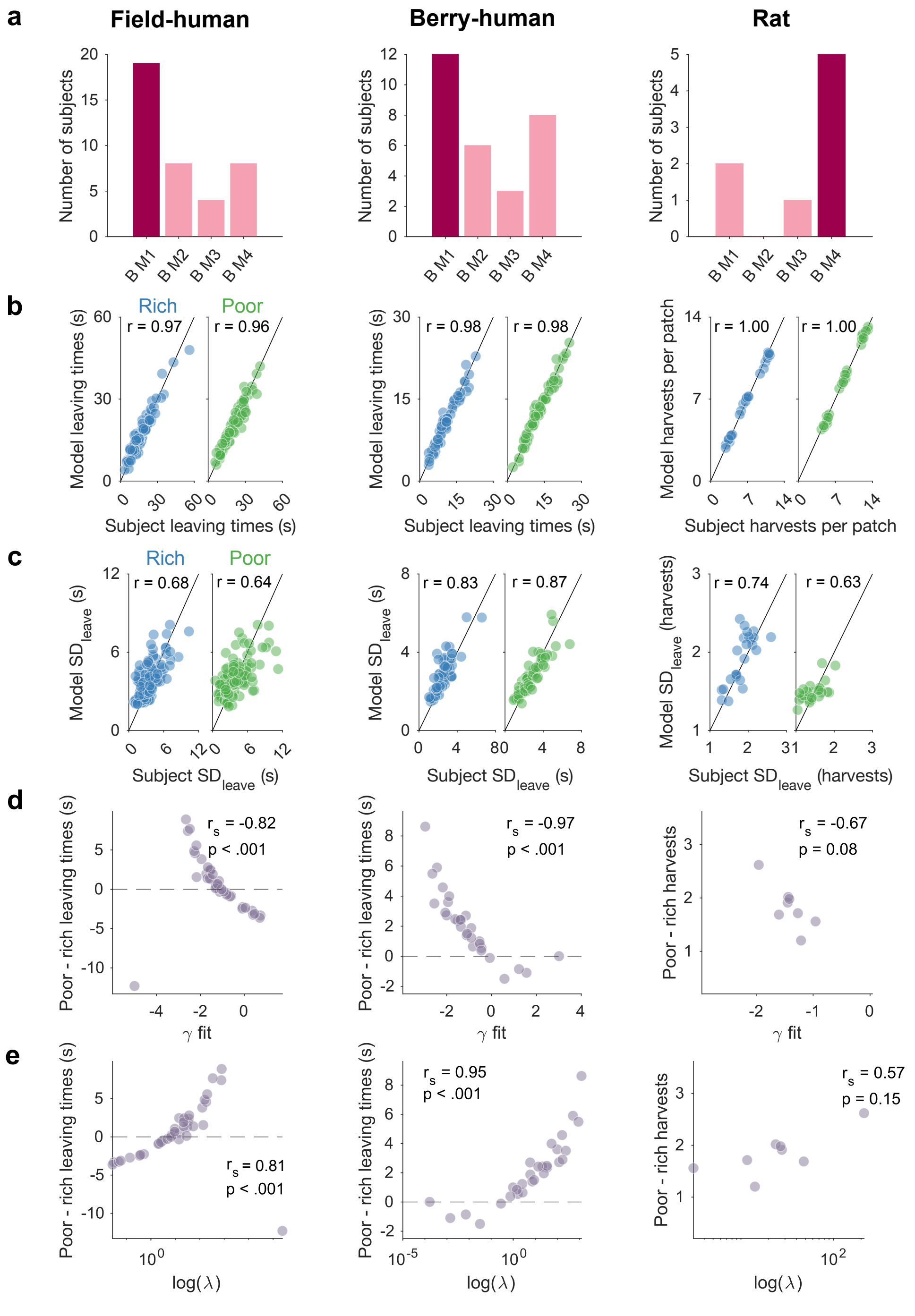


Supplementary Figure 7: **Additional model analysis for $B$ M1 - M4 a**: Number of subjects best fit by each model, according to minimum BIC. Highlighted bar shows the model best fit for the highest number of subjects. **b**: Correlation of subject and model leaving decisions. Each dot is the mean leaving time/number of harvests for each subject and patch type, separated by environment. r: Pearson's correlation coefficient. **c**: Correlation of average $SD_{leave}$ for each subject with average simulated $SD_{leave}$ for the subject. Each dot is $SD_{leave}$ for each subject and patch type, separated by environment. r: Pearson's correlation coefficients. **d**: Correlation of the magnitude of environment effect with the $\gamma$ parameter from M4 for each subject. We measured the environment effect as the difference in leaving times (or number of harvests) between poor and rich environments. Positive values indicate the subject left earlier in rich compared to poor environments. $r_s$: Spearman's correlation coefficient (field-human: n = 39, berry-human: n = 29, rat: n = 8). **e**: As per panel d, for $\lambda$ for each subject. X-axis in log-scale to better visualise dispersion of parameter values.

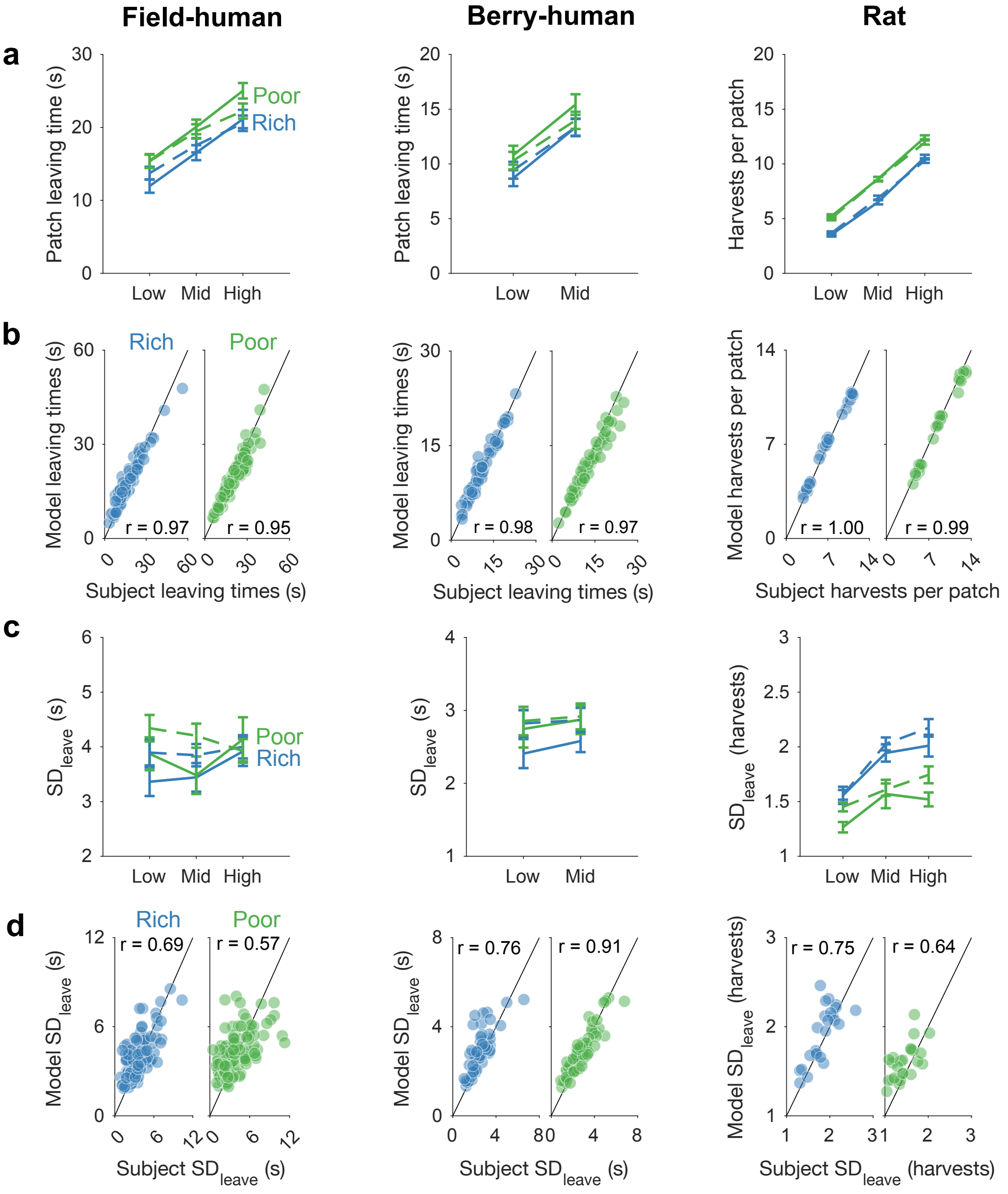


Supplementary Figure 8: **Learning variant of stochastic model can still replicate average leaving times and their variability a**: **a**: Group means of leaving times/number of harvests for the subjects (solid line) compared to the learning model (dashed line), for each patch type and environment. **b**: Correlation of subject and model leaving decisions. Each dot is the mean leaving time/number of harvests for each subject and patch type, separated by environment. r: Pearson's correlation coefficients. **c**: Group means of $SD_{leave}$ for subjects (solid line) compared to model simulations (dashed line), for each patch type and environment. **d**: Correlation of average $SD_{leave}$ for each subject with average simulated $SD_{leave}$ for the subject. Each dot is $SD_{leave}$ for each subject and patch type, separated by environment. r: Pearson's correlation coefficients.

### Supplementary Note 1: analysis of average leaving times

To examine changes in average patch leaving behaviour between environments and patch types for each task, we performed linear mixed-effects models (LMEMs) using the *lmer* function (*lme4*, v1.1-35.4) in R. LMEMs account for the within-subject design by including both fixed effects, which are assumed to remain constant across subjects, and random effects, which are assumed to vary across subjects. For example, the influence of patch or environment on leaving times is expected to vary for different subjects. Thus we included in our model both fixed and random effects of environment (rich/poor), patch type (low/mid/high) and their interaction, with a subject-level random intercept. We determined the random effects structure by first fitting the maximal model [1]. In the case of non-convergence, we reduced the random effect structure of the model based on the goals of our analysis as recommended by [1], by first removing random slopes for the interaction term (an effect of no interest), then removing random correlations, finally removing random slopes which explained no variance. We report results from models with the maximal random effects structure that converged. A priori successive difference contrasts were set for patch type and its interaction with environment, whilst environment had sum-to-zero coding. p-values were calculated using the *lmerTest* package (v3.1-3) [2] with Satterthwaite method for degrees of freedom. Confidence intervals were calculated using the profile method with the *confint* function.

We replicated the statistical results from the original articles, finding that subjects harvested high yielding patches for longer than low yielding patches (field-human: $F(2,\ 39.8) = 150$, $p = 2.2 \times 10^{-16}$; berry-human $F(1{,}27.7) = 170$, $p = 2.48 \times 10^{-13}$; rat: $F(2,\ 2586) = 3553$, $p = 2.2 \times 10^{-16}$). Subjects also harvested patches for longer in a poor environment than in a rich environment (field-human: $F(1{,}36.3) = 39.8$, $p = 2.62 \times 10^{-7}$; berry-human $F(1{,}27.7) = 26.4$, $p = 1.97 \times 10^{-5}$; rat: $F(1{,}7.4) = 163.8$, $p = 2.52 \times 10^{-6}$).

Supplementary Table 1: Linear mixed model of patch and environment predicting leaving times/number of harvests

| | *b* | *SE* | CI low | CI high | *df* | *t* | *p* |
|---|---|---|---|---|---|---|---|
| **Field-human** | | | | | | | |
| (Intercept) | 18.36 | 0.97 | 16.43 | 20.28 | 37.89 | 18.94 | < **.001** |
| Patch (mid - low) | 4.58 | 0.32 | 3.94 | 5.24 | 38.74 | 14.13 | <**.001** |
| Patch (high - mid) | 4.77 | 0.35 | 4.07 | 5.48 | 36.76 | 13.53 | <**.001** |
| Environment | 3.62 | 0.57 | 2.48 | 4.75 | 36.30 | 6.31 | <**.001** |
| Patch (mid - low) x Environment | 0.15 | 0.46 | -0.76 | 1.05 | 1876.8 | 0.32 | 0.752 |
| Patch (high - mid) x Environment | 0.46 | 0.47 | -0.45 | 1.38 | 1879.2 | 1.00 | 0.319 |
| **Berry-human** | | | | | | | |
| (Intercept) | 12.06 | 0.79 | 10.48 | 13.63 | 27.98 | 15.25 | < **.001** |
| Patch (mid - low) | 4.63 | 0.36 | 3.93 | 5.34 | 27.67 | 13.04 | <**.001** |
| Environment | 2.09 | 0.41 | 1.28 | 2.90 | 27.67 | 5.14 | <**.001** |
| Patch (mid - low) x Environment | -0.02 | 0.21 | -0.44 | 0.41 | 26.42 | -0.08 | 0.936 |
| **Rat** | | | | | | | |
| (Intercept) | 7.83 | 0.16 | 7.49 | 8.17 | 6.98 | 47.8 | < **.001** |
| Patch (mid - low) | 3.20 | 0.08 | 3.04 | 3.37 | 2589.4 | 37.9 | <**.001** |
| Patch (high - mid) | 3.93 | 0.09 | 3.76 | 4.09 | 2580.2 | 46.2 | <**.001** |
| Environment | 1.88 | 0.15 | 1.57 | 2.18 | 7.42 | 12.8 | <**.001** |
| Patch (mid - low) x Environment | 0.44 | 0.21 | 0.02 | 0.87 | 8.66 | 2.17 | 0.060 |
| Patch (high - mid) x Environment | -0.35 | 0.18 | -0.72 | 0.03 | 8.79 | -1.92 | 0.088 |

### Supplementary Note 2: analysis of variability in leaving times

To analyse patch and environment effects on variability ($SD_{leave}$), we performed a repeated measures ANOVA (Type III) using the *aov_ez* function (*afex*, v1.4-1) in R. We conducted ANOVAs instead of LMEMs as there was only a single measurement of $SD_{leave}$ per cell (each patch × environment combination). We coded the predictors using sum-to-zero coding, and conducted follow-up pairwise comparisons using *emmeans* (v1.10.2), with Tukey adjustment for multiple comparisons. Effect sizes are reported with partial eta-squared using the *effectsize* package (v1.0.0). Bayes factors for the null hypothesis, $BF_{01}$, were calculated for patch and environment using the *anovaBF* function (*BayesFactor*, v0.9.12-4.7) [3]. We used the default Cauchy prior with scale of $\frac{\sqrt{2}}{2}$. $BF_{01}$ was calculated by taking the inverse of evidence in support of the effect ($BF_{10}$). To calculate $BF_{10}$ for each main effect, we calculated the ratio of $BF_{10}$ for the full model compared to $BF_{10}$ for a restricted model in which everything except the main effect was kept. We recomputed the Bayes factors with 50,0000 iterations to reduce the proportional error on the Bayes factors below 2%.

The field-human task subjects showed no evidence for an environment effect on $SD_{leave}$ (ANOVA $F(1, 38) = 1.05$, $p$=0.31, partial eta-squared $\eta_p^2 = 0.03$; Bayes factor in favour of the null hypothesis of no effect $BF_{01} = 2.97$) and at best weak evidence for a patch effect on $SD_{leave}$ (ANOVA $F(2, 76) = 3.94$, $p$=0.024, $\eta_p^2 = 0.09$; Bayes factor for no effect $BF_{01} = 1.34$); the berry-human task subjects showed at best weak evidence for an environment effect (ANOVA $F(1,\ 28)$ = 4.08, $p$=0.053, $\eta_p^2$ = 0.13; Bayes factor $BF_{01}$ = 0.20) and no evidence for a patch effect on $SD_{leave}$ ($F(1,\ 28)$ = 2.42, $p$=0.13, $\eta_p^2$ = 0.08; Bayes factor $BF_{01}$ = 3.78).

The rats' empirical $SD_{leave}$ showed extremely strong evidence for an effect of both the type of patch (ANOVA $F(2,\ 14) = 24.46$, $p = 2.7 \times 10^{-5}$, $\eta_p^2 = 0.78$; Bayes factor for the hypothesis of an effect $BF_{10}$ = 1,365) and environment (ANOVA $F(1,\ 7) = 21.66$, $p = 0.002$, $\eta_p^2 = 0.76$; Bayes factor$BF_{10}$ = 86,633) on $SD_{leave}$.

Supplementary Table 2: Pairwise comparisons for repeated measures ANOVA: patch and environment predicting $SD_{leave}$

| | $b$ | $SE$ | CI low | CI high | $df$ | $t$ | $p$ |
|---|---|---|---|---|---|---|---|
| **Field-human** | | | | | | | |
| Patch (mid - low) | -0.16 | 0.18 | -0.60 | 0.29 | 38 | -0.86 | 0.67 |
| Patch (high - low) | 0.41 | 0.24 | -0.17 | 0.99 | 38 | 1.72 | 0.21 |
| Patch (high - mid) | 0.57 | 0.20 | 0.07 | 1.06 | 38 | 2.80 | **0.021** |
| Environment (poor - rich) | 0.26 | 0.25 | -0.25 | 0.76 | 38 | 1.03 | 0.312 |
| **Berry-human** | | | | | | | |
| Patch (mid - low) | 0.15 | 0.09 | -0.06 | 0.46 | 28 | 1.56 | 0.131 |
| Environment (poor - rich) | 0.31 | 0.16 | 0.003 | 0.53 | 28 | 2.02 | 0.053 |
| **Rat** | | | | | | | |
| Patch (mid - low) | 0.35 | 0.06 | 0.17 | 0.52 | 7 | 5.78 | **0.002** |
| Patch (high - low) | 0.35 | 0.04 | 0.23 | 0.48 | 7 | 8.58 | < **.001** |
| Patch (high - mid) | 0.008 | 0.07 | -0.19 | 0.21 | 7 | 0.12 | 0.992 |
| Environment (poor - rich) | -0.39 | 0.08 | -0.58 | -0.19 | 7 | -4.65 | **0.002** |
